# Frontal cortical regulation of neurogenesis and cellular proliferation in the ventral subventricular zone

**DOI:** 10.1101/2021.12.27.474262

**Authors:** Moawiah M Naffaa, Rehan R Khan, Chay T. Kuo, Henry H. Yin

## Abstract

Neurogenesis and differentiation of the neural stem cells (NSCs) in the subventricular zone (SVZ) are controlled by cell-intrinsic molecular pathways that interact with extrinsic signaling cues. Here we identified a novel circuit that regulates neurogenesis and cellular proliferation in the lateral ventricle SVZ (LV-SVZ). Our results demonstrate direct glutamatergic inputs from the frontal cortex as well as local inhibitory interneurons, control the activity of distinctive cholinergic neurons in the subependymal zone (subep-ChAT^+^). *In vivo* optogenetic stimulation and inhibition in this circuit were sufficient to control local SVZ neurogenesis, LV NSCs proliferation, and SVZ cellular divisions in ventral SVZ. These findings shed light on local and distal neural circuit activity-dependent regulation of postnatal and adult SVZ neurogenesis and LV-SVZ cellular proliferation.

---

The rodent subventricular zone (SVZ) of the lateral ventricles (LV) is a major site of postnatal neurogenesis ^1^. Neurogenesis in SVZ provides a useful model system for studying neuronal regeneration and tissue remodeling in the adult brain ^2,3^. LV neural stem cells (NSCs) divide asymmetrically for self-renewal or differentiate to become transient amplifying intermediate progenitors (TAPs) ^4–6^. These intermediate progenitors divide and differentiate into doublecortin-positive neuroblasts ^7,8^, which migrate to the olfactory bulb (OB) ^7,9,10^, where they become mature interneurons that are incorporated into the local circuitry ^11,12^. The generation of adult-born neurons from the SVZ contributes to experience-dependent plasticity in the postnatal brain ^13,14^ and plays a critical role in social behavior in rodents ^12,15,16^.

The LV NSCs receive inputs from a variety of sources, including neighboring NSCs, TAPs, and neuroblasts^17,18^. Different neurotransmitters, including GABA, dopamine, and serotonin, released by neurons in different brain regions, are also known to regulate postnatal and adult SVZ neurogenesis ^19–24^. However, little is known about the local and distal circuity which directly or indirectly influence the SVZ neurogenesis.

Recent work has identified a small population of cholinergic neurons in the subependymal space to the lateral ventricle. These neurons, which express Choline acetyltransferase (ChAT^+^), can modulate the proliferation of LV NSCs and neuroblasts in an activity-dependent manner ^25^. They are unlike neighboring striatal cholinergic interneurons, and release acetylcholine (ACh) into the SVZ niche. We hypothesize that the population of subependymal ChAT^+^ (subep-ChAT^+^) neurons in the ventral domain of SVZ is a key node in the neural circuit regulation of SVZ neurogenesis and LV NSCs proliferation. Using neural circuit tracing, electrophysiology and *in vivo* optogenetic strategies, we identified a novel neural circuit involving anterior cingulate glutamatergic projections and local calretinin-positive interneurons that can regulate the activity of subep-ChAT^+^ neurons. Consequently, this circuit directly modulate SVZ neurogenesis, activity of LV NSCs proliferation and SVZ cellular divisions in the ventral SVZ. These findings revealed a novel circuitry by which frontal cortical inputs can influence cholinergic signaling in ventral SVZ to modulate neurogenesis and cellular proliferation.

## Results

### Excitatory and inhibitory presynaptic inputs to the subep-ChAT^+^ neurons

Unlike striatal cholinergic neurons, subep-ChAT^+^ neurons do not show tonic activity in the absence of synaptic inputs, suggesting that excitatory inputs are needed to activate these neurons^26^. However, it is currently unknown where the excitatory inputs are generated from and how they influence the activity of subep-ChAT^+^ neurons. We hypothesized that glutamatergic inputs from cortical projection neurons may provide excitatory drive of the subep-ChAT^+^ neurons. The cortical neurons mainly express vesicular glutamate transporter 1 protein (VGlut1)^27^. To selectively stimulate these glutamatergic inputs, we crossed *VGlut1-Cre* mice with *ChAT-eGFP* and ChR2-tdTomato transgenic mouse line (*Ai27*) that expresses channelrhodopsin in a Cre-dependent manner^28^. The resulting *VGlut1-Cre; ChAT-eGFP; Ai27* mice express channelrhodopsin in all VGlut1^+^ neurons **(Fig. 1A)**.

**Figure 1.**
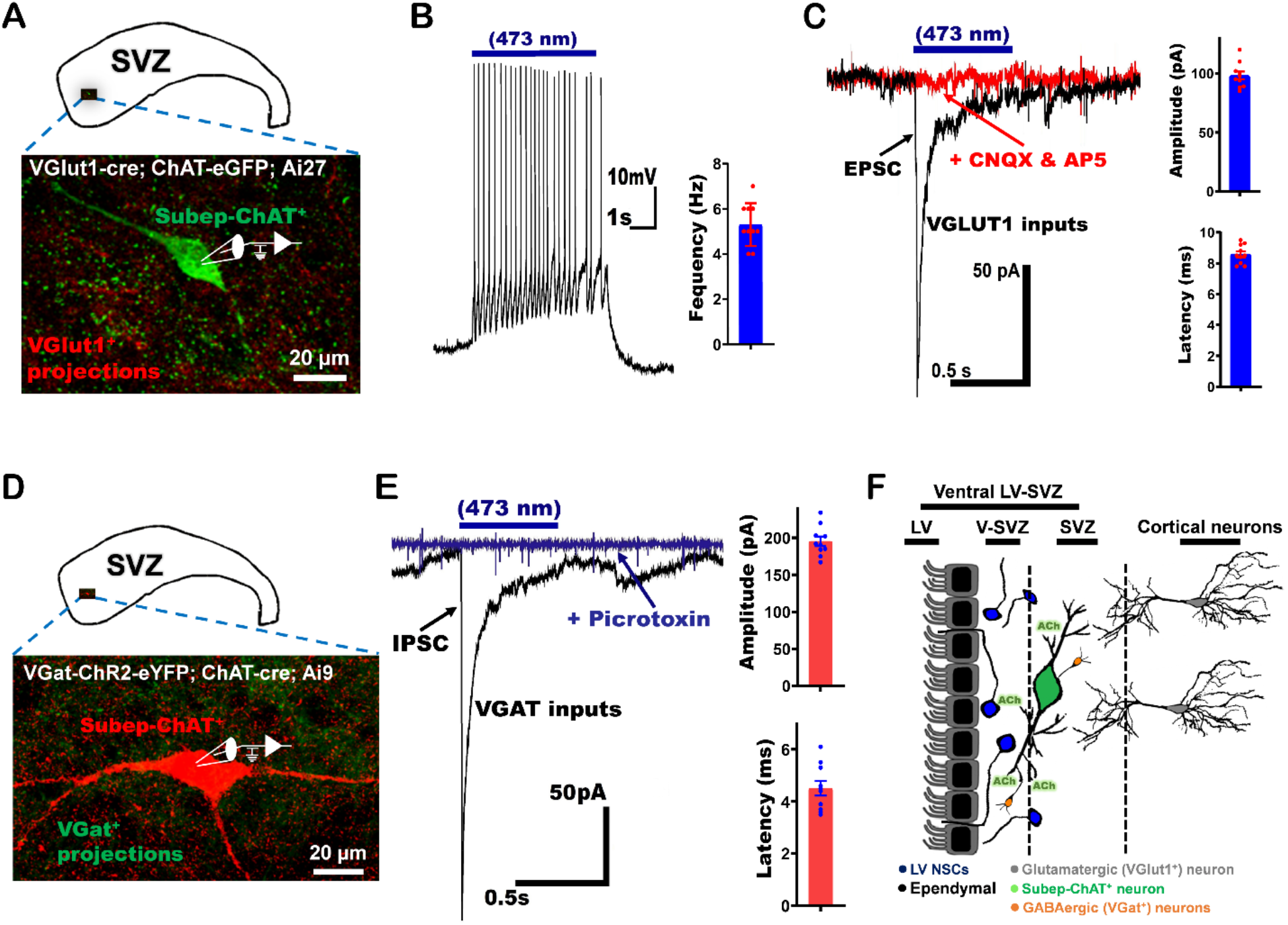
Glutamatergic and GABAergic inputs to subep-ChAT^+^ neurons. (A) Illustration of electrophysiological recording of excitatory inputs (VGlut1^+^) to the subep-ChAT^+^ neuron from the wholemount of P30-50 *VGlut1-Cre; ChAT-eGFP; Ai27* mice. (B) Representative trace from whole cell current clamp recording of evoked action potential (APs) from subep-ChAT^+^ neurons upon blue (473nm) light stimulation for 5s (Left). Blue bar = duration of light stimulation. Average frequency (Right). P < 0.0001, *t*9 = 17.7, *n* = 10. Data collected from four *VGlut1-Cre; ChAT-eGFP; Ai27* mice. Each dot represents data from one subep-ChAT^+^ neuron. (C) Representative trace of EPSCs that were obtained in whole-cell voltage-clamp recordings from subep-ChAT^+^ neurons after photostimulation for 500ms (black), and the application of AMPA and NMDA receptors antagonists; CNQX and AP-5, respectively (red) (left). Blue bar = duration of light stimulation. Average evoked EPSC amplitude and latency (Right). P < 0.0001, *t*_*9*_ = 29.1, n = 10 and P < 0.0001, *t*_*9*_ = 44.2, n = 10, respectively. Data collected from four *VGlut-Cre; ChAT-eGFP; Ai27* mice. Each dot represents the amplitude (upper) and latency (lower) of a single Excitatory Postsynaptic Current (EPSC) recorded from subep-ChAT^+^ neuron. (D) Illustration of electrophysiological recording of inhibitory inputs (VGat^+^) to the subep-ChAT^+^ neuron from P30-50 *VGat-ChR2-eYFP; ChAT-Cre; Ai9* mice. (E) Representative trace of electrophysiological recordings of IPSCs that were obtained in whole-cell recordings from subep-ChAT^+^ neurons upon blue (473nm) light stimulation for 500ms (black), and the application of GABA_A_R antagonist; picrotoxin (blue) (left). Blue bar = duration of light stimulation. Each dot represents the amplitude (upper) and latency (lower) of a single Inhibitory Postsynaptic Current (IPSC) recorded from subep-ChAT^+^ neuron. Average evoked IPSC amplitude and latency (Right). P value < 0.0001, *t*9 = 30.4, n = 10 and P value < 0.0001, *t*9 = 16.1, n = 10, respectively. Data collected from four *VGat-ChR2-eYFP; ChAT-cre; Ai9* mice. (F) Schematic representation of presynaptic (excitatory and inhibitory) inputs to the subep-ChAT^+^ neuron. All error bars indicate SEM

Whole-cell recordings from subep-ChAT^+^ neurons of *VGlut1-Cre; ChAT-eGFP; Ai27* mice showed robust firing in response to 473-nm light activation of VGlut1^+^ axon terminals (firing frequency = 5.375 ± 0.324 Hz) **(Fig. 1B)**. In voltage-clamp mode (holding at −60 mV), photo-stimulation evoked excitatory postsynaptic currents (EPSCs) in the subep-ChAT^+^ neurons. The induced currents can be blocked by glutamatergic receptors blockers AP5 (NMDA receptor antagonist) and CNQX (AMPA receptor antagonist) (Current amplitude = 98.2 ± 10.7 pA, and current latency = 8.6 ± 0.62 ms) **(Fig. 1C)**. These findings demonstrate the existence of cortical glutamatergic inputs that drive and regulate the activity of subep-ChAT^+^ neurons.

To examine the inhibitory drivers of subep-ChAT^+^ neurons, we crossed VGat-ChR2-eYFP (i.e.: channelrhodopsin expressed in GABAergic neurons) with *ChAT-Cre* and *Ai27*. The result is VGat-ChR2-eYFP; *ChAT-Cre; Ai27* mice which allow recording from subep-ChAT^+^ neurons in animals with channelrhodopsin expressed in all GABAergic neurons **(Fig. 1D)**. Whole-cell recordings from subep-ChAT^+^ neurons in voltage-clamp mode (holding at −60 mV) and activating VGAT^+^ axon terminals via 473-nm laser resulted in consistent inhibitory postsynaptic currents (IPSCs). The evoked IPSCs in the subep-ChAT^+^ neurons were blocked by picrotoxin (GABAA receptors antagonist) (Current amplitude = 196.1 ± 24.13 pA, and current latency = 4.5 ± 0.88 ms) **(Fig. 1E)**. These results suggest that subep-ChAT^+^ neurons receive direct GABAergic inputs that manage their release of ACh in the SVZ.

### Developing rabies virus strategy for tracing neural circuit

Our initial electrophysiological recording results established the presence of excitatory and inhibitory modulators to the activity of subep-ChAT^+^ neurons **(Fig. 1F)**. However, it remains unclear exactly where these inputs originated. To answer this question, we generated a Cre-dependent *R26R-FLEX-TVA-2A-RabiesG-2A-tdTomato-FLEX (R26F-RTT)* mice to trace the connectivity of subep-ChAT^+^ neurons via a single Rabies viral injection **(Fig. 2A & B)**. This new tracing tool allows for expressing TVA, RabiesG and tdTomato in all Cre-expressing neurons. The *R26F-RTT* mice were first tested by injecting EnvA G-deleted Rabies-eGFP virus into the striatum where no nonspecific rabies infections were detected at the injection site **(Extended Fig. 1A & B)**. For examining the specific expression of TVA, RabiesG and tdTomato in Cre-dependent mice, *R26F-RTT* mice were crossed with *chat-Cre, VGat-Cre and D2-Cre* to generate *R26F-RTT; chat-Cre, R26F-RTT; VGat-Cre* and *R26F-RTT; D2-Cre* mice, respectively. In *R26F-RTT; chat-Cre mice*, all ChAT^+^ neurons were only found to express tdTomato **(Extended Fig. 1C)**, which indicates that *R26F-RTT* mice are selectively expressed in Cre-positive neurons. To assess the retrogradely trans-synaptic spread of rabies with *R26F-RTT*, EnvA G-deleted Rabies-eGFP virus was injected into the striatum of *R26F-RTT; VGat-Cre* and *R26F-RTT; D2-Cre* mice **(Extended Fig. 2A & C)**. After 7-days, the cortical excitatory neurons in cingulate and secondary motor cortices which are presynaptically connected to the striatal VGat^+^ and D2^+^ neurons were found labeled with GFP **(Extended Fig. 2B & D)**. This indicates that rabies can retrogradely spread from Cre-positive VGat^+^ and D2^+^ neurons when they are crossed with *R26F-RTT* mice. Together, these results demonstrated that *R26F-RTT* mice can express TVA, RabiesG and tdTomato selectively in Cre-expressing neurons and allows for retrograde presynaptic neuronal tracing.

**Figure 2.**
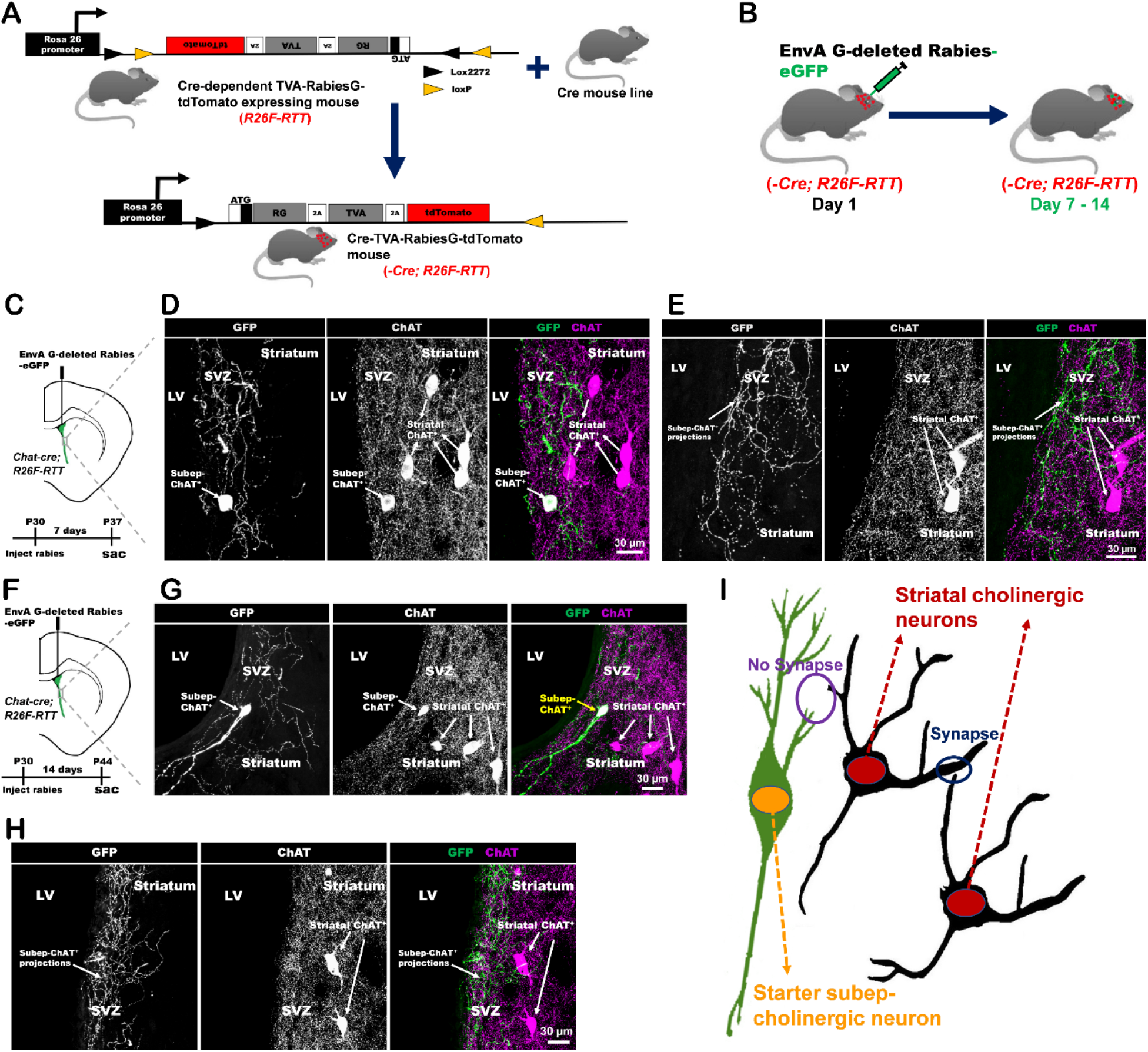
Rabies virus infection of subep-ChAT^+^ neurons. (A) Schematic representation of *R26R-FLEX-TVA-2A-RabiesG-2A-tdTomato-FLEX* (R26F-RTT) mice before and after Cre recombinase. (B) Schematic of EnvA G-deleted Rabies-eGFP virus injection into the brain of *-Cre; R26F-RTT* mice. (C) Experimental representation of EnvA G-deleted Rabies-eGFP virus injection into lateral ventricle (LV) of P30 *Chat-Cre; R26F-RTT* mice. Mice were sacrificed on the day 7 after injection. (D-E) Immunofluorescence staining for GFP (green) and ChAT (red) in SVZ and striatum of ipsilateral (injected) side of *Chat-Cre; R26F-RTT* mice (7 days post-injection). (F) Experimental representation of EnvA G-deleted Rabies-eGFP virus injection into LV of P30 *Chat-Cre; R26F-RTT* mice. Mice were sacrificed on the day 14 after injection. (G-H) Immunofluorescence staining for GFP (green) and ChAT (red) in SVZ and striatum of the ipsilateral (injected) side of *Chat-Cre; R26F-RTT* mice (14 days post-injection). (I) Schematic representation of intra cholinergic connections of striatal cholinergic and subep-ChAT^+^ neurons.

To allow for neural tracing of cholinergic neurons, we generated *ChAT-Cre; R26F-RTT* mice to express TVA receptor, RabiesG protein, and tdTomato in all cholinergic neurons **(Fig. 2B)**. We first injected EnvA G-deleted rabies-eGFP virus into the striatum of P30 *ChAT-Cre; R26F-RTT* mice **(Extended Fig. 3A)**. As the striatal cholinergic neurons receive synaptic inputs from each other^29^, an infection of a striatal cholinergic neuron with EnvA G-deleted Rabies-eGFP virus results in polysynaptic labeling of other neighbor striatal cholinergic neurons, which subsequently pass rabies to other connected neurons **(Extended Fig. 3B & C)**. Then, EnvA G-deleted Rabies-eGFP virus was injected into the LV of *ChAT-Cre; R26F-RTT* mice to target subep-ChAT^+^ neurons, immediately following mannitol injection to disrupt the ependymal layer locally **(Fig. 2C & F)**. Using this strategy, we were able to selectively infect a subep-ChAT^+^ neuron and trace their presynaptic inputs **(Fig. 2D & G)**. The local connectivity of subep-ChAT^+^ neurons and their projections were checked at 7- and 14-days post rabies injection **(Fig. 2D, E, G & H)**. Unlike striatal cholinergic neurons, subep-ChAT^+^ neurons in the SVZ niche are not connected with neighboring striatal cholinergic neurons **(Fig. 2I)**. These results confirm that subep-ChAT^+^ neurons have distinct neuronal connectivity.

### Local presynaptic inhibitory input to subep-ChAT^+^ neurons

Injection of EnvA G-deleted Rabies-eGFP virus into the LV of *ChAT-Cre; R26F-RTT* mice revealed local GFP^+^ interneurons adjacent to the infected subep-ChAT^+^ neurons **(Fig. 3A & B)**. While staining for markers of the known inhibitory neurons, colocalization between GFP^+^ interneurons and calretinin (CR) antibody was noticed **(Fig. 3C)**.

**Figure 3.**
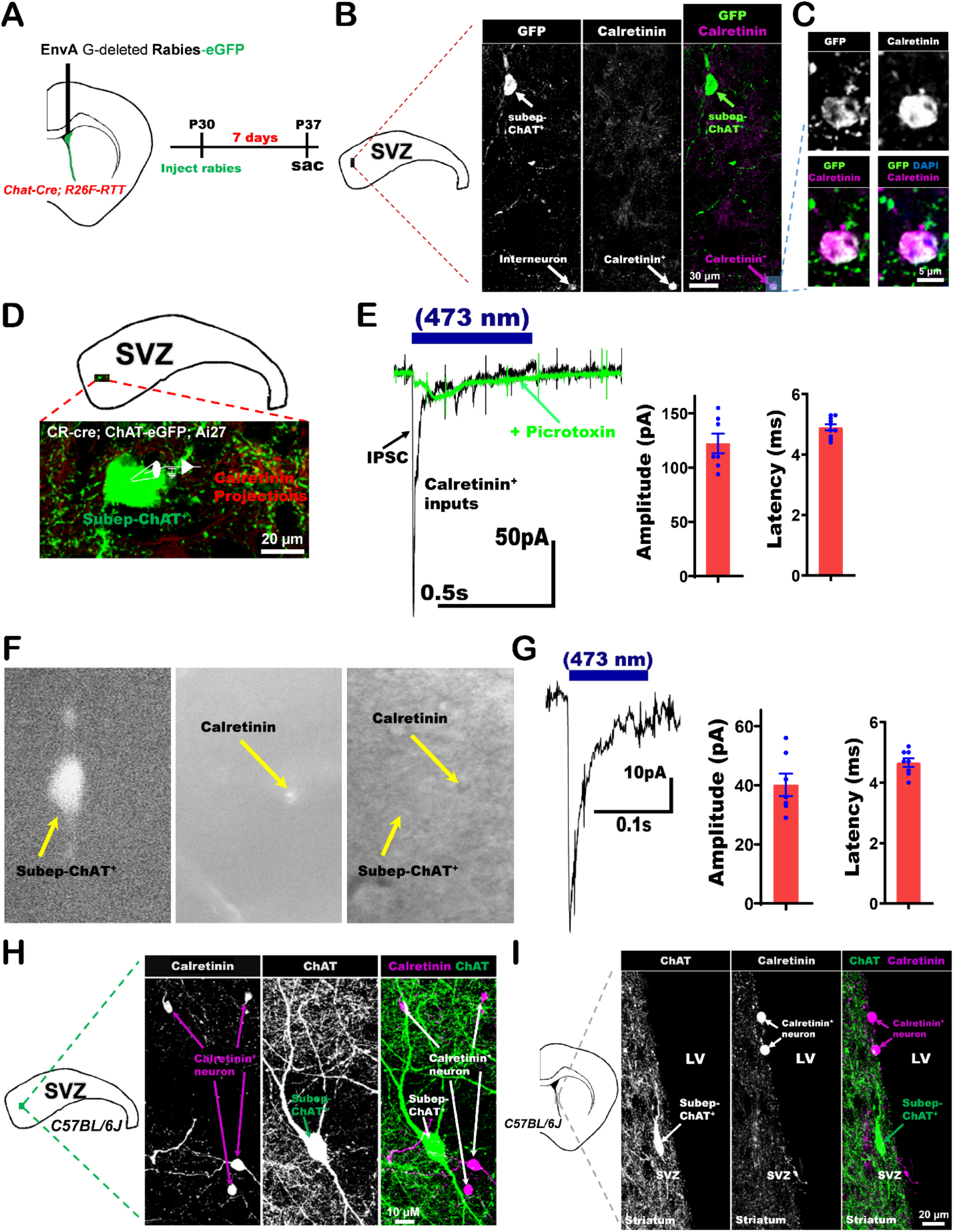
Local calretinin-positive (CR^+^) GABAergic interneurons provide inhibitory inputs to subep-ChAT^+^ neurons. (A) Experimental design of EnvA G-deleted Rabies-eGFP virus injection into LV of P30 *Chat-Cre; R26F-RTT* mice. (B-C) Immunofluorescence staining for GFP (green), Calretinin (purple) and DAPI (blue) in ipsilateral SVZ wholemount (injected) of P37 *Chat-Cre; R26F-RTT* mice. (D) illustration of electrophysiological recording of calretinin-positive (CR^+^) inhibitory inputs to the subep-ChAT^+^ neurons from SVZ wholemount of P30-50 *Cr-Cre; ChAT-eGFP; Ai27* mice. (E) Representative trace of evoked IPSCs from subep-ChAT^+^ neurons upon photostimulation for 500ms (black) and after application of GABAAR antagonist picrotoxin (green) (Left). Blue bar = duration of light stimulation. Average current amplitude and latency of the IPSCs upon stimulation (Right). P < 0.0001, *t*_*9*_ = 19.1, n = 10 and P < 0.0001, *t*_*9*_ = 46.9, n = 10, respectively. Data collected from five *Cr-Cre; ChAT-eGFP; Ai27* mice. Each dot represents the amplitude (left) and latency (right) of a single Inhibitory Postsynaptic Current (EPSC) recorded from subep-ChAT^+^ neuron. (F) Representation of subep-ChAT^+^ neuron recording from SVZ wholemount of P30-50 *Cr-Cre; ChAT-eGFP; Ai27* mice. Left; subep-ChAT^+^ neuron (473nm light). Middle; Subep-CR^+^ neurons (590nm light). Right; bright field image showing both subep-ChAT^+^ and -CR^+^ neurons. (G) Representative trace of evoked IPSCs in subep-ChAT^+^ neurons following photostimulation for 100ms (Left). Blue bar = duration of light stimulation. Average current amplitude and latency of the IPSCs upon stimulation (Right). P < 0.0001, *t*_*7*_ = 12.25, n = 8 and P < 0.0001, *t*_*9*_ = 32.4, n = 8, respectively. Data collected from three *Cr-Cre; ChAT-eGFP; Ai27* mice. Each dot represents the amplitude (left) and latency (right) of an Inhibitory Postsynaptic Current (EPSC) recorded from subep-ChAT^+^ neuron. (H-I) Immunofluorescence staining for CR (purple) and ChAT (green) in the SVZ wholemount and coronal section (I) of P28 *C57BL/6J* mice. All error bars indicate SEM.

To test if the CR^+^ interneurons provide local inhibitory inputs to subep-ChAT^+^ neurons, we performed whole-cell patch clamp recording from subep-ChAT^+^ neurons of *Cr-Cre; ChAT-eGFP; Ai29* mice **(Fig. 3D)**. A high-chloride internal solution was used to detect inhibitory postsynaptic currents (IPSCs) while holding at −60 mV in voltage clamp. We stimulated local CR^+^ interneurons with blue light and revealed robust evoked IPSCs, which were completely blocked by picrotoxin (current amplitude = 122.3 ± 9.0 pA, and current latency = 4.9 ± 0.33 ms) **(Fig. 3E)**. Then, we tested for other known inhibitory interneurons expressing calbindin (CB^+^), somatostatin (SST^+^) or parvalbumin (PV^+^) **(Extended Fig. 4A)**. Using whole-cell recording from subep-ChAT^+^ neurons, an optogenetic stimulation of GABA released from these inhibitory interneurons (i.e.: CB, SST and PV) did not induce IPSCs **(Extended Fig. 4B)**.

To further confirm the local CR^+^ interneuron inputs to the subep-ChAT^+^ neurons, we used a Patterned Light Stimulator LED controller (Mightex) to target CR^+^ labelled neurons in SVZ wholemount of *Cr-Cre; ChAT-eGFP; Ai29* mice. Focal optogenetic activation of a single CR^+^ interneuron generated IPSCs in subep-ChAT^+^ neuron (current amplitude = 40.14 ± 3.78 pA, and current latency = 4.7 ± 0.4 ms) **(Fig. 3F & G)**. Subsequently, we stained against CR and ChAT using *C57BL/6J* mice, revealing the presence of 2-4 CR^+^ interneurons surrounding each subep-ChAT^+^ neuron in the LV-SVZ **(Fig. 3H & I)**. Together these results confirm that local CR^+^ interneurons are the main source of inhibitory inputs to the subep-ChAT^+^ neuron.

### Presynaptic excitatory inputs from cingulate cortex area 1 (Cg1) to subep-ChAT^+^ neurons

To illuminate long distance connections for subep-ChAT^+^ neurons in the brain, EnvA rVSV-eGFP (EnvA/RABVG-eGFP) virus^30^ was injected into the LV of *ChAT-Cre; R26F-RTT* mice **(Fig. 4A)**. Distal GFP^+^ projection neurons from the anterior cingulate cortex area 1 (Cg1) were observed in the ipsilateral hemisphere only **(Fig. 4B)**. To confirm the Cg1 input to subep-ChAT^+^ neurons, pAAV-CaMKIIa-hChR2(E123A)-mCherry virus was first injected into the Cg1 region of *C57BL/6J* mice to express mCherry in the GFP^+^ neurons from Figure 4B **(Fig. 4C)**. By tracing their projections in the SVZ region, we observed them adjacent to the subep-ChAT^+^ neurons **(Fig. 4D)**.

**Figure 4.**
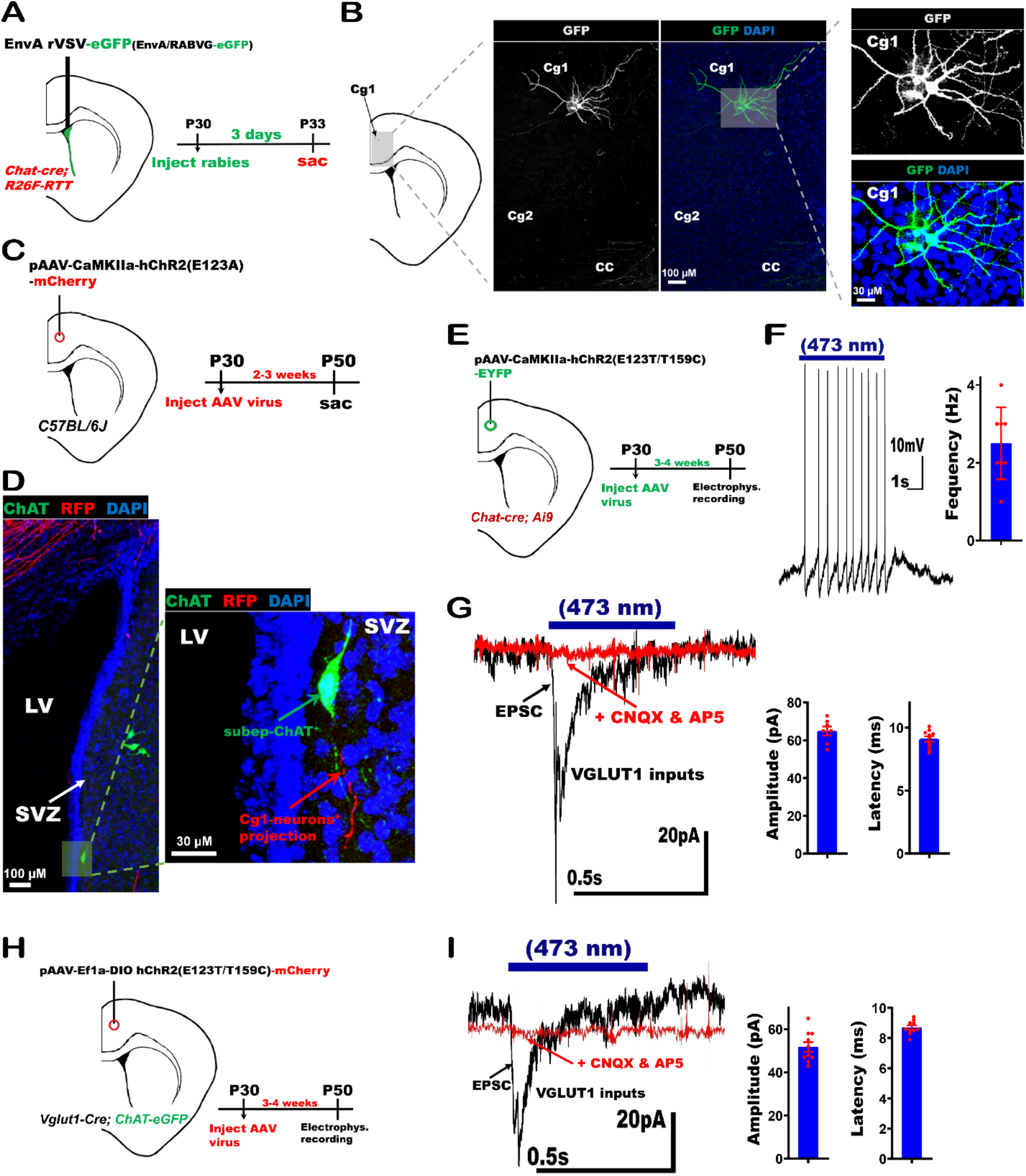
Anterior cingulate cortex projects directly to subep-ChAT^+^ neurons. (A) Schematic illustration of EnvA rVSV-eGFP (EnvA/RABVG-eGFP) virus injection into LV of P30 *Chat-Cre; R26F-RTT* mice. (B) A sample of immunofluorescence staining for GFP (green) in anterior cingulate cortex areas 1 and 2 (Cg1 and Cg2) (left) and Cg1 region (right) of ipsilateral (injected) side of P33 *Chat-Cre; R26F-RTT* mice. (C) Schematic representation of AAV-CaMKII-hChR2(E123A)-mCherry virus injection into Cg1 region of P30 *C57BL/6J* mice. (D) A sample of immunofluorescence staining for ChAT (green) and infected projections (red) of the ipsilateral Cg1 neurons (injected side) in striatum and SVZ (left), and SVZ (right) of P50 *C57BL/6J* mice in panel C. (E) Experimental design of pAAV-Ef1a-DIO hChR2(E123T/T159C)-EYFP virus injection into Cg1 region of P30 *Chat-Cre; Ai9* mice. Mice were used for recording 3-4 weeks post-injection. (F) Representative trace of whole cell current clamp recoding of evoked action potentials (APs) from subep-ChAT^+^ neurons upon photo-stimulation for 5s (Left). Blue bar = duration of light stimulation. Average spike frequency (Right). P < 0.0001, *t*_*7*_ = 7.6, n = 8. Data collected from five *Chat-Cre; Ai9* mice. Each dot represents data from one subep-ChAT^+^ neuron. (G) Representative trace of EPSCs were obtained in whole-cell recordings from subep-ChAT^+^ neurons upon 473nm light stimulation for 500ms (black), and after blocking with AMPA and NMDA receptors antagonists (CNQX and AP-5, respectively) (red) (Left). Blue bar = duration of light stimulation. Average EPSC amplitude and latency (Right). P value < 0.0001, *t*9 = 36.16, n = 10 and P value < 0.0001, *t*9 = 40.5, n = 10, respectively. Data collected from five *Chat-Cre; Ai9* mice. Each dot represents the amplitude (left) and latency (right) of a single Excitatory Postsynaptic Current (EPSC) recorded from subep-ChAT^+^ neuron. (H) Experimental design of pAAV-Ef1a-DIO hChR2(E123T/T159C)-mCherry virus injection into Cg1 region of P30 *VGlut1-Cre; ChAT-eGFP* mice. Mice were used for recording at 3-4 weeks post-injection. (I) Representative trace of EPSCs were obtained in whole-cell voltage-clamp recordings from subep-ChAT^+^ neurons upon 473nm light stimulation for 500ms (black), and after blocking with AMPA and NMDA receptors antagonists (CNQX and AP-5, respectively) (red) (Left). Blue bar = duration of light stimulation. Average evoked EPSC amplitude and latency (Right). P value < 0.0001, *t*9 = 23.3, n = 10 and P value < 0.0001, *t*9 = 61.2, n = 10, respectively. Data collected from four *VGlut1-Cre; ChAT-eGFP* mice. Each dot represents the amplitude (left) and latency (right) of a single Excitatory Post Synaptic Current (EPSC) recorded from subep-ChAT^+^ neuron. All error bars indicate SEM.

To functionally examines a monosynaptic input from Cg1 to the subep-ChAT^+^ neurons, a general ChR2 viral vector (pAAV-CaMKIIa-hChR2(E123T/T159C) -EYFP) was injected into the Cg1 region of *Chat-Cre; Ai9* mice to express ChR2 in the projections of Cg1 neurons **(Fig. 4E)**. Optogenetic stimulation of the cingulate cortex projections reliably evoked action potentials in subep-ChAT^+^ neurons (firing frequency = 2.5 ± 0.327 Hz) **(Fig. 4F)**. Using voltage clamp, the optogenetic stimulation induced reliable EPSCs in subep-ChAT^+^ neurons which were completely blocked by the glutamate antagonists (AP5 and CNQX) (current amplitude = 64.86 ± 2.424 pA, and current latency = 9.1 ± 0.7 ms) **(Fig. 4G)**. These results confirm the presence of glutamatergic neurons in the ipsilateral Cg1 region that excite subep-ChAT^+^ neurons.

To further confirm the (Cg1-subep-ChAT^+^) circuit, pAAV-Ef1a-DIO hChR2(E123T/T159C)-mCherry virus was injected into the Cg1 region of *VGlut1-Cre; ChAT-eGFP* mice to express ChR2 in Cg1 VGlut1^+^ neurons **(Fig. 4H)**. In the voltage clamp mode, reliably EPSCs were generated in subep-ChAT^+^ neurons upon optogenetic stimulation of Cg1 VGlut1^+^ neurons projection **(Fig. 4I, Left)**. The induced EPS currents were completely blocked by AP5 and CNQX, confirming the presence of Cg1 glutamatergic inputs (current amplitude = 53.29 ± 2.96 pA, and current latency = 8.7 ± 0.44 ms) **(Fig. 4I, Right)**. These findings show that a specific population of cortical neurons in the Cg1 region provides excitatory drive to subep-ChAT^+^ neurons.

### *Calretinin-Cre (Cr-Cre)* mice label the distal Cg1 excitatory presynaptic input to the subep-ChAT^+^ neurons

After identifying Cg1 glutamatergic input into subep-ChAT^+^ neurons using rabies neural tracing and electrophysiological recording, (Cg1-subep-ChAT^+^) circuit was then further tested by injecting AAVrg viruses (i.e.: retrograde tracing using AAV viruses) into LVs of *C57BL/6J, VGlut1-Cre* and *CR-Cre* mice. Upon injection of pAAVrg-hSyn-mCherry and pAAVrg-hSyn-DIO-mCherry viruses into LVs of *C57BL/6J* and *VGlut1-Cre* mice, respectively **(Extended Fig. 5A & C)**, we observed mCherry^+^ neurons in the ipsilateral Cg1 region **(Extended Fig. 5B & D)**. Surprisingly, injecting pAAVrg-hSyn-DIO-mCherry virus into the LV of *CR-Cre* mice labels neurons in the ipsilateral Cg1 region **(Fig. 5A &B)**. The detected mCherry^+^ neurons in the Cg1 of *C57BL/6J, VGlut1-Cre* and *CR-Cre* mice post pAAVrg virus injections and the GFP^+^ Cg1 neurons in **Fig. 4B** are located along the same rostro-caudal axis of the Cg1 region. These findings strongly suggest that the Cg1 presynaptic glutamatergic drivers of subep-ChAT^+^ neurons are calretinin^+^ (CR^+^) during the postnatal period.

**Figure 5.**
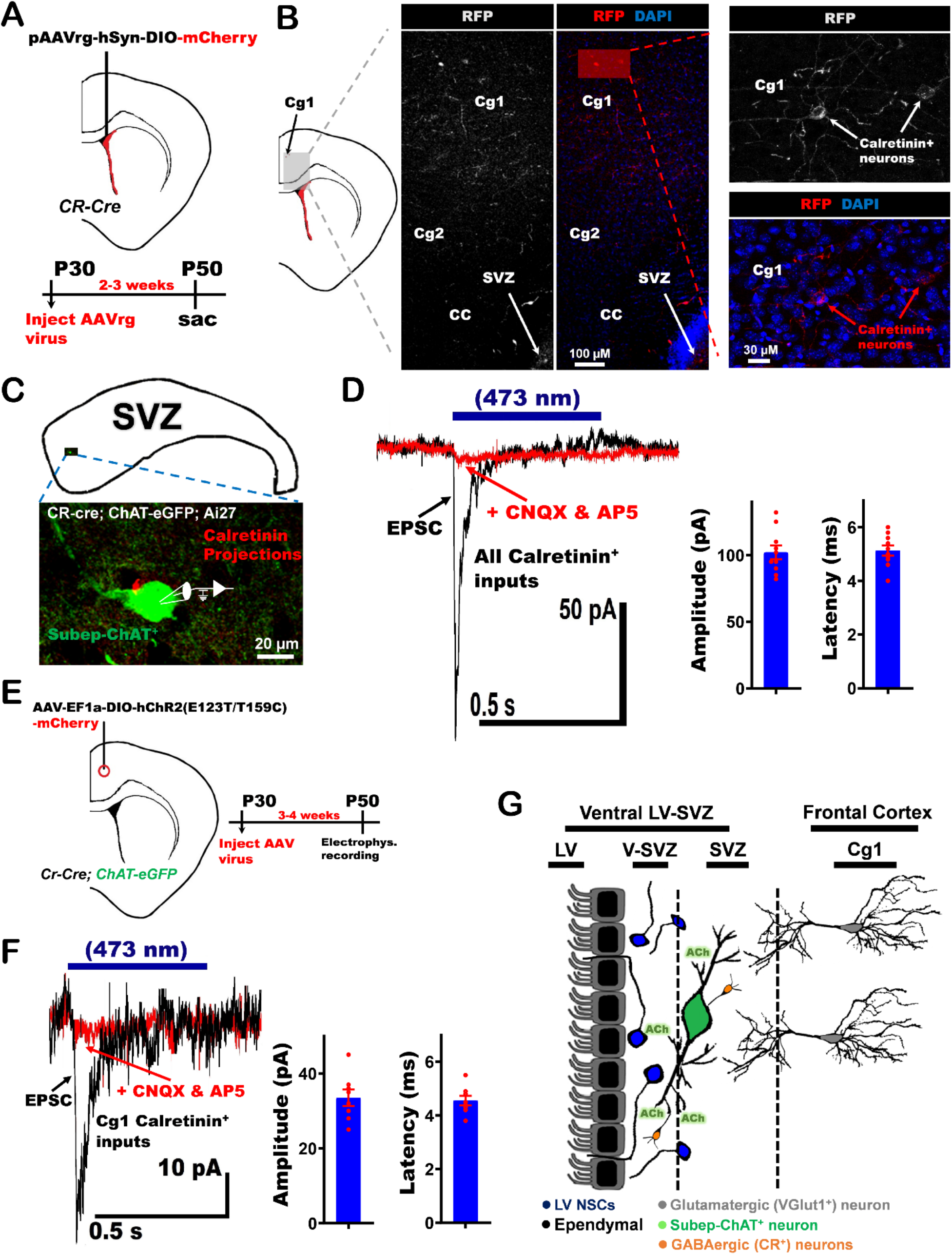
Presynaptic cingulate neurons that excite subep-ChAT^+^ neurons are calretinin-positive (CR^+^). (A) Schematic illustration of the experimental design of pAAVrg-hSyn-DIO-mCherry virus injection into LV of P28 *Cr-Cre* mice. (B) Immunofluorescence staining for RFP (red) in ipsilateral (injected side) anterior cingulate cortex areas 1 and 2 (Cg1 and Cg2) (left) and ipsilateral (injected side) Cg1 region (right) of P50 *Cr-Cre* mice. (C) Illustration of electrophysiological recording of calretinin-positive (CR^+^) excitatory inputs to subep-ChAT^+^ neurons from SVZ wholemount of P30-50 *Cr-Cre; ChAT-eGFP; Ai27* mice. (D) Representative trace of EPSCs were obtained in whole-cell recordings from subep-ChAT^+^ neurons upon 473nm light stimulation for 500ms (black), and after blocking with AMPA and NMDA receptors antagonists (CNQX and AP-5, respectively) (red) (Left). Blue bar = duration of light stimulation. Average EPSC amplitude and latency (Right). P < 0.0001, *t*_*9*_ = 19.6, n = 10 and P < 0.0001, *t*_*9*_ = 27, n = 10, respectively. Data collected from five *Cr-Cre; ChAT-eGFP; Ai27* mice. Each dot represents the amplitude (left) and latency (right) of a single Excitatory Postsynaptic Current (EPSC) recorded from subep-ChAT^+^ neuron. (E) Experimental design of AAV-EF1a-DIO-hChR2(E123T/T159C)-mCherry virus injection into Cg1 of P28 *Cr-Cre; ChAT-eGFP* mice. Mice were used for recording 3-4 weeks post-injection. (F) Representative trace of EPSCs evoked specifically by Cg1 CR^+^ input. EPSCs were recorded from subep-ChAT^+^ neurons of injected mice in panel E upon 473nm light stimulation for 500ms (black). EPSCs were blocked by CNQX and AP-5 (red) (Left). An average current amplitude and latency (Right). P < 0.0001, *t*_*7*_ = 15.3, n = 8 and P < 0.0001, *t*_*7*_ = 24.9, n = 8, respectively. Data collected from four *Cr-Cre; ChAT-eGFP* mice. Each dot represents the amplitude (left) and latency (right) of a single Excitatory Postsynaptic Current (EPSC) recorded from subep-ChAT^+^ neuron. (G) Schematic summary of (Cg1-Subep-ChAT^+^) circuit regulation of LV NSCs. All error bars indicate SEM.

To examine that these Cg1 glutamatergic inputs to the subep-ChAT^+^ neurons are CR^+^ neurons, whole-cell recordings from subep-ChAT^+^ neurons were first performed in *Cr-Cre; ChAT-eGFP; Ai27* mice **(Fig. 5C)**. In voltage mode at −60 mV, optogenetic stimulation of CR^+^ terminals reliably evoked EPSCs (Current amplitude = 106 ± 6.68 pA, and current latency = 5.13 ± 0.44 ms), which were completely blocked by AP5 and CNQX **(Fig. 5D)**. This predicts that the population of Cg1 glutamatergic that drives the activity of subep-ChAT^+^ neurons are CR^+^ postnatally. To further functionally testing whether these Cg1 excitatory CR^+^ neurons regulate the activity of subep-ChAT^+^ neurons, we injected AAV-EF1a-DIO-hChR2(E123T/T159C)-mCherry virus into the Cg1 region of *Cr-Cre; ChAT-eGFP* mice **(Fig. 5E)**. Whole-cell recording from subep-ChAT^+^ neurons in voltage clamp mode induced EPSCs upon optogenetic stimulation (Current amplitude = 33 ± 2.46 pA, and latency = 5.13 ± 0.44 ms) **(Fig. 5F)**. All previous data confirm that a specific population of cortical neurons in Cg1 region (VGlut1^+^ and CR^+^) provides excitatory drive to subep-ChAT^+^ neurons **(Fig. 5G)**.

The total number of labeled Cg1 cortical neurons with *Cr-Cre* mice during postnatal period is must lower than the number of labeled neurons with *VGlut1-Cre* mice. Thereby, the *Cr-Cre* mouse is a reliable tool to study the population of glutamatergic neurons from the Cg1 region that control the activity of subep-ChAT^+^ neurons.

### *in vivo* (Cg1-subep-ChAT^+^) circuit stimulation regulate neurogenesis in the ventral domain of SVZ

Based on our earlier findings, we defined the Cg1 presynaptic excitatory drivers of subep-ChAT^+^ neurons. Thereafter, we studied the *in vivo* functional role of (Cg1-subep-ChAT^+^) circuit on SVZ neurogenesis and LV cellular proliferation. We hypothesized that this circuit regulate SVZ neurogenesis surrounding subep-ChAT^+^ neurons by modulating the cellular proliferation in the ventral SVZ. To fulfill this aim, AAV-EF1a-DIO-hChR2 (E123T/T159C)-mCherry virus was injected into ipsilateral Cg1, and optical fibers were implanted into the Cg1 regions of *Cr-Cre* mice **(Fig. 6A)**. The optogenetic stimulation was continuously conducted for 2- or 3-days and delivered by TTL control of 473-nm laser, 10 ms pulses at 10 Hz, lasting 10 s, given once every 1 min **(Fig. 6B and Extended Fig. 6A)**.

**Figure 6.**
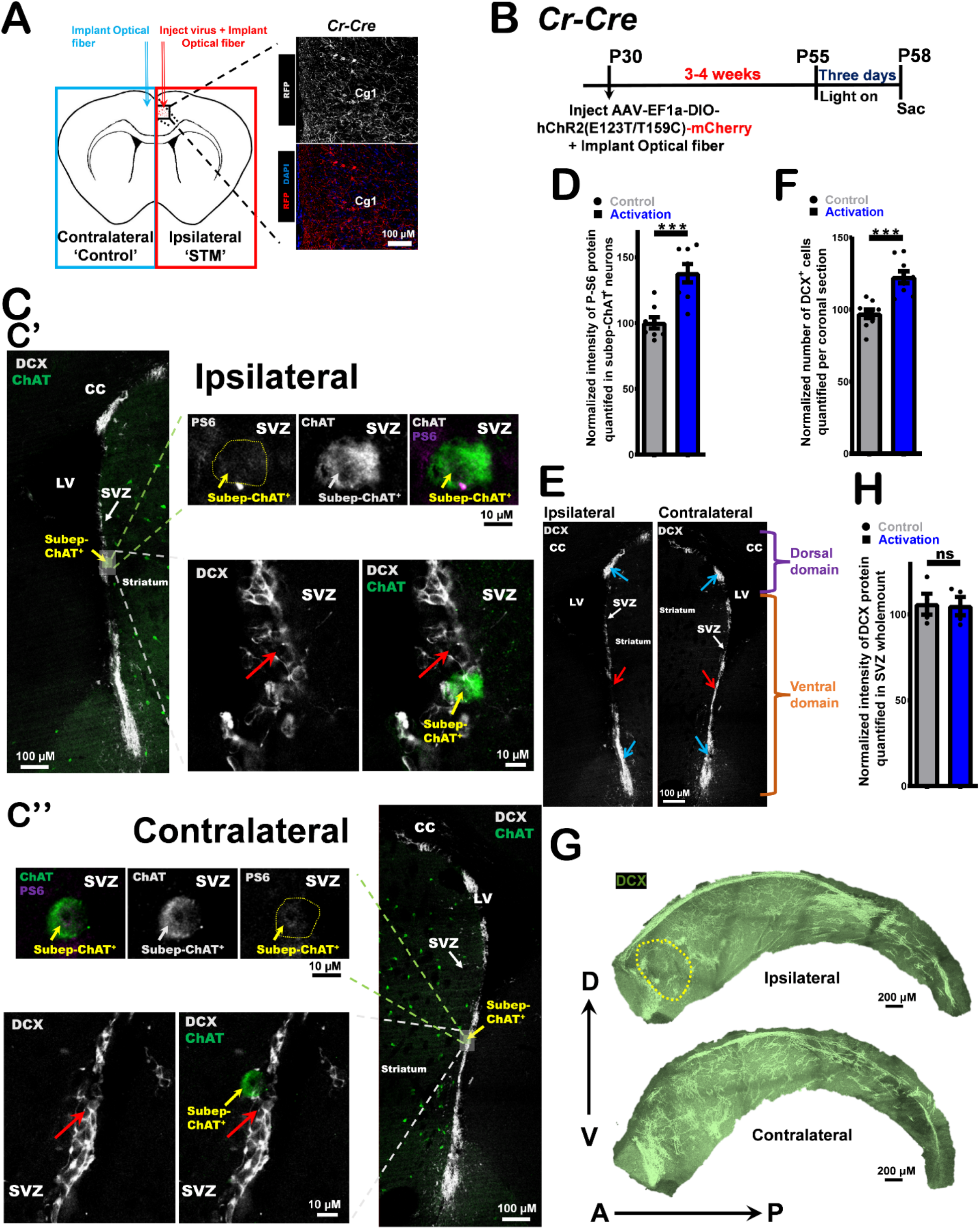
*In vivo* optogenetic (Cg1-subep-ChAT^+^) circuit stimulation modulates neurogenesis in the SVZ niche. (A) Schematic representation of *in vivo* optogenetic stimulation experiment in panel B. Immunofluorescence staining for RFP (red) in the ipsilateral (injected) Cg1 of *Cr-Cre* mice. (B) Experimental design of *in vivo* optogenetic stimulation for three days post AAV-EF1a-DIO-hChR2(E123T/T159C)-mCherry virus injection into ipsilateral Cg1 and optical fibers implantation into Cg1 regions of P28 *Cr-Cre* mice (C) DCX (grey) and ChAT (green) immunofluorescence staining of (C’) ipsilateral (upper images; activation) vs. (C’’) contralateral (lower images; control) sides of SVZ from stimulated coronal sections of mice in panel B. Identical settings from same brain section were used for imaging. Red arrows represent DCX^+^ neuroblasts around subep-ChAT^+^ neurons. (D) P-S6 intensity analysis of subep-ChAT^+^ neurons in ipsilateral side (activation) vs. contralateral side (control). P<0.0004, n=8, Paired t-test. Data collected from three stimulated *Cr-Cre* mice. Each dot represents a subep-ChAT^+^ neuron. (E) DCX (grey) immunofluorescence staining of ipsilateral SVZ (left image; activation) vs. contralateral SVZ (right image; control) from stimulated coronal section of mice in panel C. Identical settings from same brain section were used for imaging. Blue arrows display DCX^+^ neuroblasts in dorsal (upper) and ventral (lower) domains. Red arrow displays DCX^+^ neuroblasts adjacent to subep-ChAT^+^ neurons in ventral domain of the stimulated SVZ (left) and control SVZ (right). (F) Analysis of DCX^+^ neuroblasts in stimulated coronal sections of ipsilateral side (activation) vs. contralateral side (control). P<0.0001, n=9, Paired t-test. Data collected from four stimulated *Cr-Cre* mice. Each dot represents a total DCX^+^ cells per coronal section. (G) DCX (green) immunofluorescence staining of ipsilateral (upper images; activation) vs. contralateral (lower images; control) of SVZ wholemount from stimulated mice in panel B. Identical settings from same brain wholemounts were used for imaging. Yellow dotted circle represents an area where subep-ChAT^+^ neurons mainly reside in the ventral region of SVZ. (H) Analysis of DCX intensity in ipsilateral SVZ wholemount (activation) vs. contralateral SVZ wholemount (control). P = 0.766 (ns), n=4, Paired t-test. Data collected from four stimulated *Cr-Cre* mice. Each dot represents a total DCX protein per SVZ wholemount. All error bars indicate SEM.

In the 2-days circuit stimulation experiments, the mice were perfused immediately after stimulation cessation and brain slices of the stimulated Cg1 sections were stained against ChAT, phospho-S6 ribosomal (P-S6) and doublecortin (DCX) antibodies. To confirm (Cg1-subep-ChAT^+^) circuit activation in the studied coronal sections, the neuronal activity of subep-ChAT^+^ neurons in these sections was determined by quantifying the intensity of P-S6 protein **(Extended Fig. 6B (B’ & B’’); upper)**. Among the included coronal sections in our studies, subep-ChAT^+^ neurons on the ipsilateral side showed significantly higher P-6S intensity than their counterpart on the contralateral side within the same brain (43.9 ± 7.3%) **(Extended Fig. 6C)**. In these sections, the doublecortin (DCX^+^) neuroblasts surrounding subep-ChAT^+^ neurons on the activation side were not clustered and were mainly disordered unlike their counterpart DCX^+^ neuroblasts on the contralateral side **(Extended Fig. 6B (B’ & B’’); lower)**. This observed disorganized pattern of the clustered local DCX^+^ cells adjacent to the activated subep-ChAT^+^ neurons may be related to the continuous stimulation of the circuit which result in a local constant release of ACh. Although the circuit was stimulated for two days, the quantification of the total number of DCX^+^ neuroblasts showed no significant difference between the ipsilateral (activated) and contralateral (control) sides of these coronal sections **(Extended Fig. 6D)**. The total DCX^+^ cells in the whole SVZ were also assessed by quantifying the intensity of DCX staining on the ipsilateral SVZ (stimulated) vs. contralateral SVZ (control) **(Extended Fig. 6E)**. The intensity of DCX was remarkably higher in the activated SVZ compared to the control SVZ (17.7 ± 6.6%) **(Extended Fig. 6F)**.

In (Cg1-subep-ChAT^+^) circuit stimulation experiment for 3-days, the neuronal activity of subep-ChAT^+^ neurons was first examined in the coronal sections which were used for functional studies **(Fig. 6C (C’ & C’’); upper)**. These sections have shown remarkable higher P-S6 intensity in the ipsilateral (activated) subep-ChAT^+^ neurons compared to the subep-ChAT^+^ neuron in the contralateral (control) side of the same brain (37.8 ± 8.2%) **(Fig. 6D)**. The DCX^+^ neuroblasts surrounding the subep-ChAT^+^ neurons on the activation side were highly randomized and not clustered as their comparative DCX^+^ cells on the contralateral side **(Fig. 6C (C’ & C’’); lower (red arrows))**. Due to the persistent stimulation of subep-ChAT^+^ neurons, constant generation of DCX^+^ neuroblasts was observed, thus many neuroblasts migrate before they cluster together **(Fig. 6E; red arrows)**. Notably, the higher rate of newly produced DCX^+^ neuroblasts in the activated side compared to the control side led to more accumulation of neuroblasts on the margins of the dorsal and ventral SVZ domains **(Fig. 6E; blue arrows)**. The total number of DCX^+^ cells in the stimulated coronal section of the ipsilateral side was significantly higher than on the contralateral side (25.2 ± 4.3%) **(Fig. 6F)**. Although, the intensity of DCX staining of the stimulated and control SVZ wholemounts does not appear to be significantly different from each other **(Fig. 6G & H)**, it was observed to be less in the area surrounding subep-ChAT^+^ neurons of the stimulated ventral SVZ compared to the control ventral SVZ **(Fig. 6G; yellow dotted circle)**.

### *in vivo* (Cg1-subep-ChAT^+^) circuit stimulation regulate cellular proliferation in the ventral domain of SVZ

So far, we demonstrated the role of (Cg1-subep-ChAT^+^) circuit in regulating the activity of ventral SVZ neurogenesis. Subsequently, we studied the impact of this circuit on the regional LV cellular proliferation around subep-ChAT^+^ neurons in the ventral SVZ. To achieve this goal, AAV-EF1a-DIO-hChR2 (E123T/T159C)-mCherry virus was injected into ipsilateral Cg1, and optical fibers were implanted into the Cg1 regions of *Cr-Cre* mice **(Fig. 7A; right)**. To test the stimulation effect of (Cg1-subep-ChAT^+^) circuit on the LV-SVZ cellular proliferation, the circuit was activated for one day and 5-Ethynyl-2′-deoxyuridine (EdU) was intraperitoneally (IP) injected within the last 2-3 hours before mice were sacrificed. **(Fig. 7A; left)**. Due to the spatial cellular arrangements of proliferative cells and subep-ChAT^+^ neurons in the LV-SVZ neurogenic niche, we used SVZ wholemount to observe the cellular proliferation in the V-SVZ (i.e.: the proliferative cells underneath ependymal cells and 4-6um above subep-ChAT^+^ neurons) and SVZ (i.e.: the proliferative cells lining around the subep-ChAT^+^ neurons) **(Fig. 7B)**.

**Figure 7.**
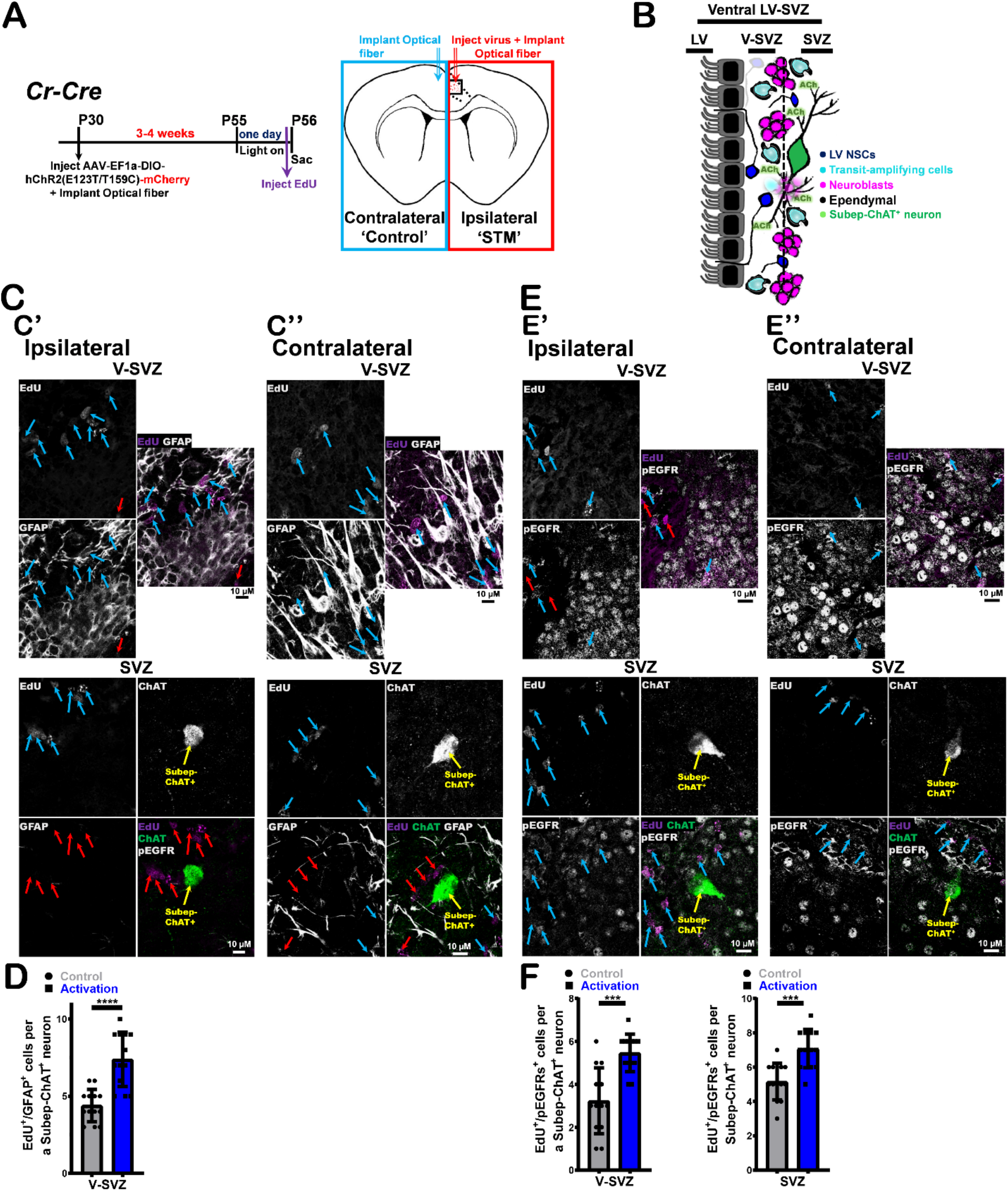
*In vivo* optogenetic (Cg1-subep-ChAT^+^) circuit stimulation modulates cellular proliferation in the SVZ niche. (A) Experimental design of *in vivo* optogenetic stimulation for one day post AAV-EF1a-DIO-hChR2(E123T/T159C)-mCherry virus injection into ipsilateral Cg1 and optical fibers implantation into Cg1 regions of P28 *Cr-Cre* mice (B) Schematic representation of the cellular composition and organization of the LV-SVZ. (C) EdU (purple), ChAT (green) and GFAP (grey) immunofluorescence staining of (C’) ipsilateral SVZ wholemount (activation) (upper images; V-SVZ and lower images; SVZ) vs. (C’’) contralateral SVZ wholemount (control) (upper images; V-SVZ and lower images; SVZ) from stimulated mice in panel A. Identical settings from same brain section were used for imaging. Blue arrows display EdU^+^/GFAP^+^ cells surrounding subep-ChAT^+^ neurons in V-SVZ. Red arrows display EdU^+^/GFAP^-^ cells surrounding subep-ChAT^+^ neurons in V-SVZ and SVZ. (D) Analysis of EdU^+^/GFAP^+^ cells in ipsilateral (activation) V-SVZ vs. contralateral (control) V-SVZ of SVZ wholemounts. P<0.0001, n=13, Unpaired t-test. Data collected from four stimulated *Cr-Cre* mice. Each dot represents total EdU^+^/GFAP^+^ cells surrounding a subep-ChAT^+^ neuron. (E) EdU (purple), ChAT (green) and pEGFR (grey) immunofluorescence staining of (E’) ipsilateral SVZ wholemount (activation) (upper images; V-SVZ and lower images; SVZ) vs. (E’’) contralateral SVZ wholemount (control) (upper images; V-SVZ and lower images; SVZ) from stimulated mice in panel A. Identical settings from same brain section were used for imaging. Blue arrows display EdU^+^/pEGFR^+^ cells surrounding subep-ChAT^+^ neurons in V-SVZ and SVZ. Red arrows display EdU^+^/pEGFR^-^ cells surrounding subep-ChAT^+^ neurons in V-SVZ. (F) Left: Analysis of EdU^+^/pEGFR^+^ cells in ipsilateral (activation) V-SVZ vs. contralateral (control) V-SVZ of SVZ wholemounts. P = 0.0001, n=13, Unpaired t-test. Data collected from four stimulated *Cr-Cre* mice. Each dot represents total EdU^+^/pEGFR^+^ cells surrounding a subep-ChAT^+^ neuron. Right: Analysis of EdU^+^/pEGFR^+^ cells in ipsilateral (activation) SVZ vs. contralateral (control) SVZ of SVZ wholemounts. P = 0.0002, n=13, Unpaired t-test. Data collected from four stimulated *Cr-Cre* mice. Each dot represents total EdU^+^/pEGFR^+^ cells surrounding a subep-ChAT^+^ neuron. All error bars indicate SEM.

Prior to observing the cellular proliferation in the area around subep-ChAT^+^ neurons, we first examined the neuronal activity of subep-ChAT^+^ neurons in the stimulated SVZ wholemount compared to their counterpart in the control SVZ wholemount. A mouse in each group of the stimulated mice was used to inspect and analyze the neuronal activity of subep-ChAT^+^ neurons **(Extended Fig. 7A)**. In these devoted mice, the activity of subep-ChAT^+^ neurons in the stimulated SVZ was higher than in the control ones (31.6 ± 7.8%) **(Extended Fig. 7B)**. A combination of EdU with either GFAP (glial fibrillary acidic protein) or pEGFR (epidermal growth factor receptor (phospho Y1068)) markers were used to view the cellular proliferation adjacent to subep-ChAT^+^ neurons in the ventral SVZ **(Fig. 7C & E)**. While GFAP^+^ cells (i.e.: A marker of quiescent NSCs (qNSCs) and early activated NSCs (aNSCs))^6^ were observed mainly in the V-SVZ **(Fig. 7C)**, the number of spatially localized EdU^+^/GFAP^+^ cells above the subep-ChAT^+^ neurons in the ipsilateral V-SVZ (activated) **(Fig. 7C’)** were higher than in the contralateral V-SVZ (control) **(Fig. 7C’’)** (3 ± 0.56%) **(Fig. 7D)**. This shows that (Cg1-subep-ChAT^+^) circuit regulates the proliferative activity of LV NSCs in the ventral SVZ.

pEGFR is a known marker for labeling both the aNSCs and transit amplifying cells (TAC)^6^ which was utilized here to study the cellular proliferation in the V-SVZ and SVZ post (Cg1-subep-ChAT^+^) circuit stimulation **(Fig. E)**. In the ipsilateral V-SVZ, the number of spatially localized EdU^+^/pEGFR^+^ cells above the subep-ChAT^+^ neurons were notably higher than in the control V-SVZ **(Fig. 7E’ & E’’; upper)** (2.23 ± 0.5%) **(Fig. 7F; left)**. This strongly suggests that (Cg1-subep-ChAT^+^) circuit has a role in the regulation of LV NSCs activity. In addition, there are more observed EdU^+^/pEGFR^+^ cells around subep-ChAT^+^ neurons in the stimulated SVZ than in control SVZ **(Fig. 7E’ & E’’; lower)** (1.92 ± 0.43%) **(Fig. 7F; right)**. The presence of more proliferative cells in both the V-SVZ and SVZ following the (Cg1-subep-ChAT^+^) circuit stimulation proposes that this circuit is involved in modulating the activity of NSCs and cellular division in the LV-SVZ.

### *in vivo* (Cg1-subep-ChAT^+^) circuit inhibition modulate SVZ neurogenesis and LV cellular proliferation in the ventral SVZ

Our previous results predicted that the (Cg1-subep-ChAT^+^) circuit controls neurogenesis and cellular proliferative activity in the ventral domain of LV-SVZ. To further understand the role of this circuit, we hypothesized that *in vivo* optogenetic inhibition of (Cg1-subep-ChAT^+^) circuit is sufficient to modulate the activity of SVZ neurogenesis around subep-ChAT^+^ neurons. To achieve that, pAAV_hSyn1-SIO-stGtACR2-FusionRed virus was injected into ipsilateral Cg1, and optical fibers were implanted into the Cg1 regions of *Cr-Cre* mice **(Fig. 8A)**. The inhibition of Cg1 CR^+^ neurons were continuously conducted for two days and were delivered by TTL control of a 473-nm laser **(Fig. 8B)**. The mice were perfused directly after the circuit inhibition was terminated and brain slices of the targeted sections were stained against DCX, ChAT and P-S6 antibodies. The coronal sections included in our studies showed higher neuronal activity of subep-ChAT^+^ neurons on the ipsilateral (inhibited) side compared to the activity of subep-ChAT^+^ neurons on the control side within the same mice brain **(Fig. 8C (C’ & C’’); upper)** (−31.25 ± 9.03%) **(Fig. 8D)**. Adjacent to the subep-ChAT^+^ neurons on the ipsilateral (inhibited) side, the number of DCX^+^ neuroblasts were significantly lower than their counterpart on the control side **(Fig. 8C (C’ & C’’); lower)**. These results indicate that the (Cg1-subep-ChAT^+^) circuit regulates local neurogenesis in the ventral SVZ. Furthermore, the total number of DCX^+^ neuroblasts on the ipsilateral side was lower than their comparative neuroblasts on the contralateral ones (−16.4 ± 3.2%) **(Fig. 8E & F)**. All previous findings proposed a crucial role for (Cg1-subep-ChAT^+^) circuit in managing neurogenesis of areas surrounding subep-ChAT^+^ neurons in the ventral SVZ.

**Figure 8.**
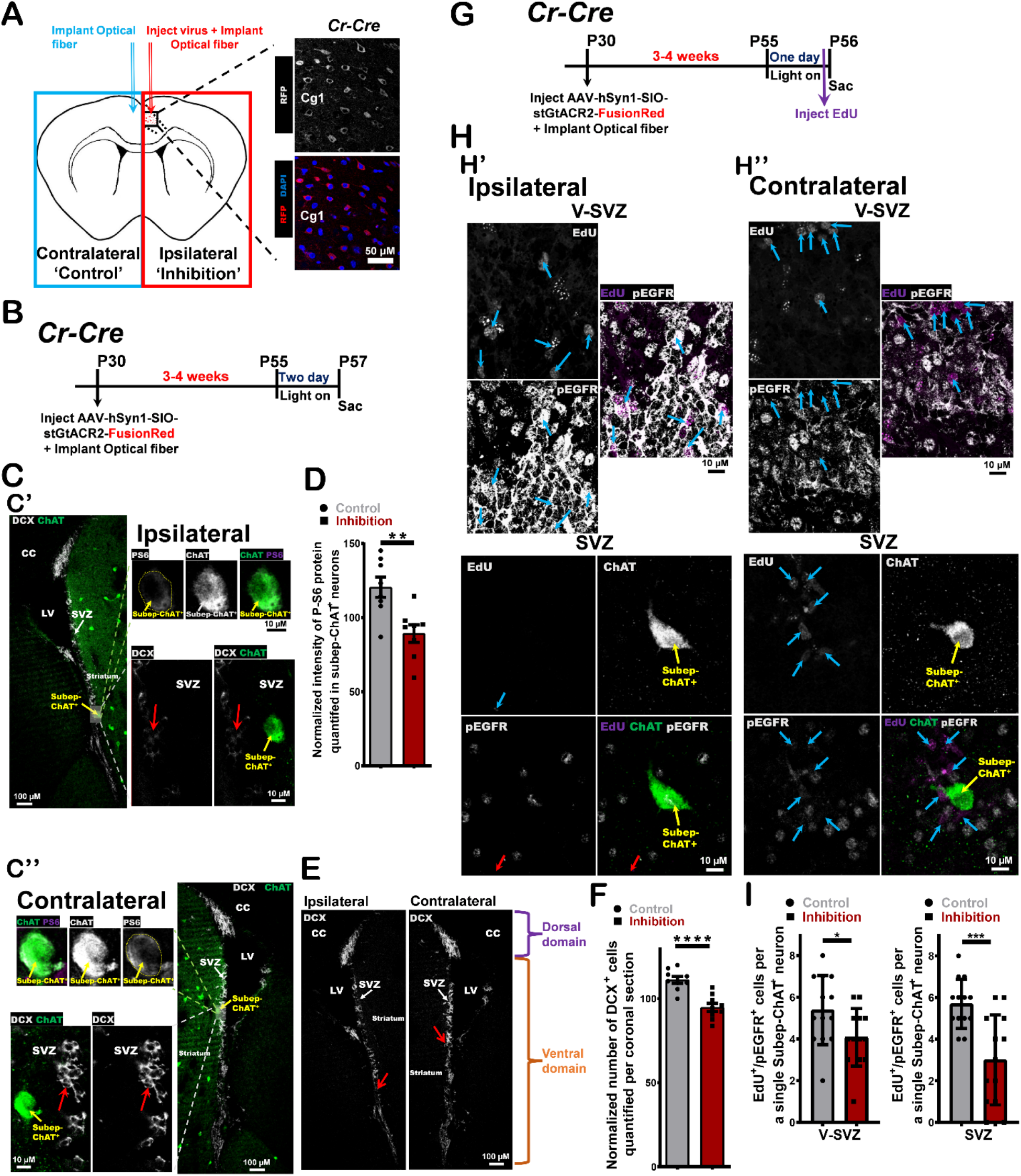
*In vivo* optogenetic (Cg1-subep-ChAT^+^) circuit inhibition modulates neurogenesis and cellular proliferation in the SVZ niche. (A) Schematic representation of *in vivo* optogenetic inhibition experiment in panel B. Immunofluorescence staining for RFP (red) in ipsilateral (injected) Cg1 of *Cr-Cre* mice. (B) Experimental design of *in vivo* optogenetic inhibition for two days post AAV_hSyn1-SIO-stGtACR2-FusionRed virus injection into ipsilateral Cg1 and optical fibers implantation into Cg1 regions of P28 *Cr-Cre* mice. (C) DCX (grey) and ChAT (green) immunofluorescence staining of (C’) ipsilateral (upper images; inhibition) vs. (C’’) contralateral (lower images; control) sides of SVZ from stimulated coronal sections of mice in panel B. Identical settings from same brain section were used for imaging. Red arrows represent DCX^+^ neuroblasts around subep-ChAT^+^ neurons. (D) P-S6 intensity analysis of subep-ChAT^+^ neurons in ipsilateral side (inhibition) vs. contralateral side (control). P = 0.0038, n=8, Paired t-test. Data collected from three stimulated *Cr-Cre* mice. Each dot represents a Subep-ChAT^+^ neuron. (E) DCX (grey) immunofluorescence staining of ipsilateral SVZ (left image; inhibition) vs. contralateral SVZ (right image; control) from stimulated coronal section of mice in panel C. Identical settings from same brain section were used for imaging. Red arrow displays DCX^+^ cells adjacent to subep-ChAT^+^ neurons in ventral domain of the stimulated and control sides. (F) Analysis of DCX^+^ cells in stimulated coronal sections of ipsilateral side (inhibition) vs. contralateral side (control). P<0.0001, n=9, Paired t-test. Data collected from four stimulated *Cr-Cre* mice. Each dot represents a total DCX^+^ cells per coronal section. (G) Experimental design of *in vivo* optogenetic inhibition for one day post AAV_hSyn1-SIO-stGtACR2-FusionRed virus injection into ipsilateral Cg1 and optical fiber implantation into Cg1 regions of P28 *Cr-Cre* mice. (H) EdU (purple), ChAT (green) and pEGFR (grey) immunofluorescence staining of (H’) ipsilateral SVZ wholemount (inhibition) (upper images; V-SVZ and lower images; SVZ) vs. (H’’) contralateral SVZ wholemount (control) (upper images; V-SVZ and lower images; SVZ) from stimulated mice in panel G. Identical settings from same brain section were used for imaging. Blue arrows display EdU^+^/pEGFR^+^ cells surrounding subep-ChAT^+^ neurons in V-SVZ and SVZ. Red arrows display EdU^+^/pEGFR^-^ cells surrounding subep-ChAT^+^ neurons in V-SVZ and SVZ. (I) Left: Analysis of EdU^+^/pEGFR^+^ cells in ipsilateral (inhibition) V-SVZ vs. contralateral (control) V-SVZ of SVZ wholemounts. P = 0.039, n=13, Unpaired t-test. Data collected from four stimulated *Cr-Cre* mice. Each dot represents the total EdU^+^/pEGFR^+^ cells surrounding a subep-ChAT^+^ neuron. Right: Analysis of EdU^+^/pEGFR^+^ cells in ipsilateral (inhibition) SVZ vs. contralateral (control) SVZ of SVZ wholemounts. P = 0.0006, n=13, Unpaired t-test. Data collected from four stimulated *Cr-Cre* mice. Each dot represents total EdU^+^/pEGFR^+^ cells surrounding a subep-ChAT^+^ neuron. All error bars indicate SEM.

Subsequently, the inhibition effect of (Cg1-subep-ChAT^+^) circuit on the LV-SVZ cellular proliferative activity around subep-ChAT^+^ neurons was investigated. To accomplish this, pAAV_hSyn1-SIO-stGtACR2-FusionRed virus was injected into ipsilateral Cg1, and optical fibers were implanted into the Cg1 regions of *Cr-Cre* mice **(Fig. 8A)**. The (Cg1-subep-ChAT^+^) circuit was continuously blocked for one day and EdU injected within the last 2-3 hours before mice were sacrificed **(Fig. 8G)**. The neuronal activity of subep-ChAT^+^ neurons was examined in the ipsilateral SVZ and compared to their counterpart in control SVZ within every studied group of mice as described in the previous section **(Extended Fig. 8A)**. In the used mice for this purpose, the subep-ChAT^+^ neurons in the ipsilateral SVZ wholemounts have shown remarkably lower activity than their contralateral counterparts (−38.1 ± 7.57%) **(Extended Fig. 8B)**. Upon staining for GFAP and EdU markers, the number of spatially localized EdU^+^/GFAP^+^ cells above subep-ChAT^+^ neurons in the ipsilateral V-SVZ **(Extended Fig. 9A’)** was noticeably less than in the control V-SVZ **(Extended Fig. 9A’’)** (−1.77 ± 0.55%) **(Fig. 9B)**. This suggests that (Cg1-subep-ChAT^+^) circuit modulates the proliferative activity of LV-NSCs. Furthermore, both pEGFR and EdU markers were applied together to study the effect of circuit inhibition on the proliferative activity of LV aNSCs and TAC in the V-SVZ and SVZ, respectively. The number of EdU^+^/pEGFR^+^ cells that spatially resides over the subep-ChAT^+^ neurons in the ipsilateral V-SVZ was observably lower than in the contralateral V-SVZ **(Fig. 8H’ & H’’; upper)** (−2.5 ± 0.1%) **(Fig. 8I; left)**. Additionally, the noticed number of EdU^+^/pEGFR^+^ cells around subep-ChAT^+^ neurons in the ipsilateral SVZ was less than in control SVZ **(Fig. 8H’ & H’’; lower)** (−2.7 ± 0.68%) **(Fig. 8I; right)**. Together, these findings suggest that (Cg1-subep-ChAT^+^) circuit has regulatory roles on the proliferative activity of NSCs and cellular division in the ventral LV-SVZ.

**Figure 9.**
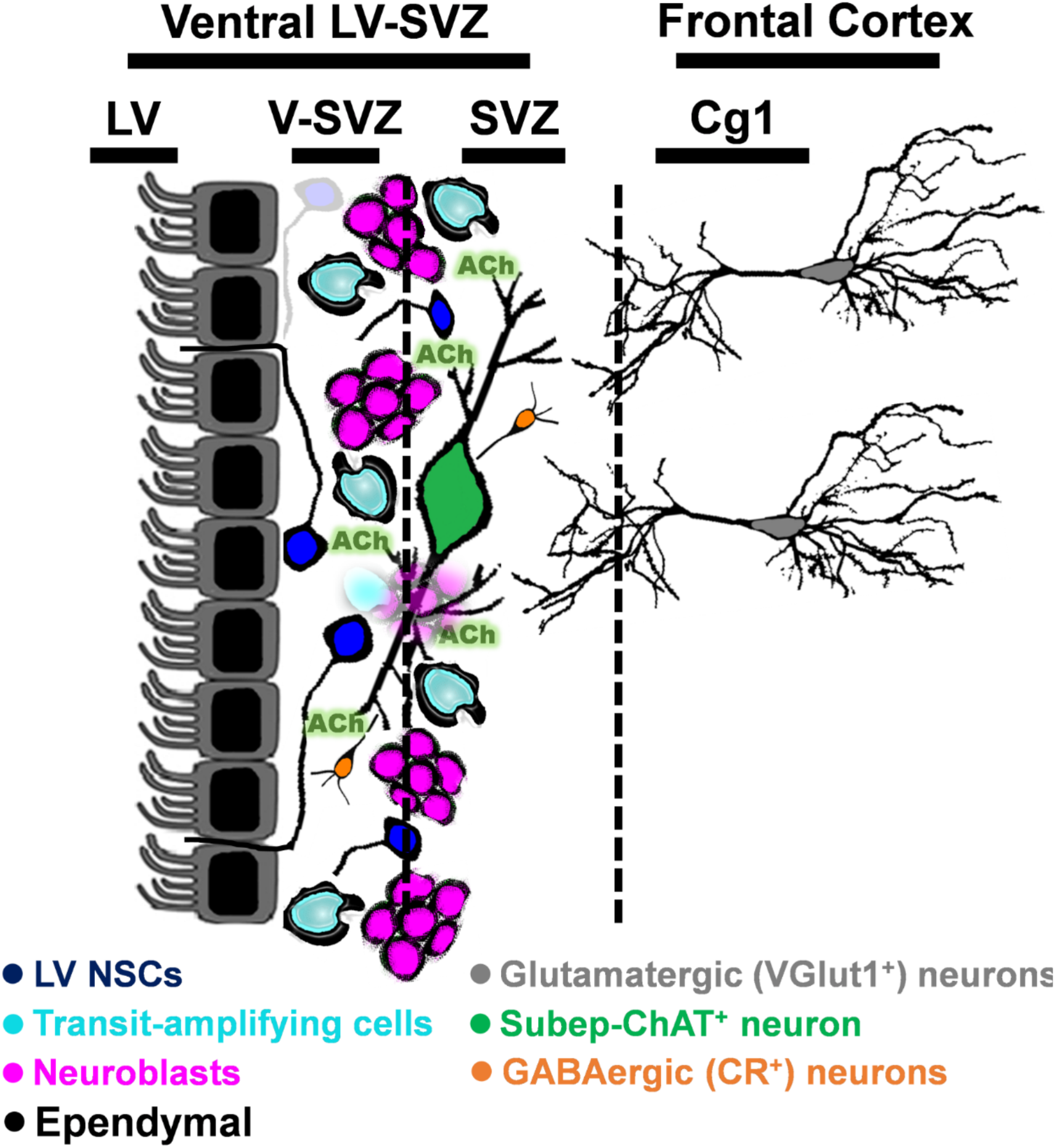
Schematic summary of presynaptic inputs into subep-ChAT^+^ neuron within LV-SVZ niche.

## Discussion

Postnatal and adult LV NSCs proliferation and SVZ neurogenesis are known to be modulated by neural activity ^18^. Here, we identified a novel circuit controlling SVZ neurogenesis and LV-SVZ cell proliferation in the LV-SVZ. Using a Rabies tracing strategy, we determined the source of presynaptic inputs to the subep-ChAT^+^ neurons, which are demonstrated to direct LV NSCs proliferation^25^. Our results showed that subep-ChAT^+^ neurons have a different pattern of neural connectivity than the striatal cholinergic neurons which are connected with their adjacent striatal cholinergic neurons^31^. Interestingly, we found that glutamatergic inputs from a specific VGlut1^+^ neuronal population in the anterior cingulate cortex area 1 (Cg1) that projects directly to subep-ChAT^+^ neurons. This is the first identification of a distal cortical input that monosynaptically drives subep-ChAT^+^ neurons and regulate their activity in the SVZ niche. In addition, we also identified a small population of local GABAergic calretinin (CR^+^) interneurons that directly inhibit subep-ChAT^+^ neurons. Previously, GABA was shown to be involved in various functions in the SVZ such as enhancing neuroblast maturation ^32^, and preserving the postnatal/adult LV NSCs by inhibiting their proliferation and differentiation, but the source of GABA was largely unclear ^21,33^. We demonstrated here for the first time GABAergic CR^+^ interneurons are a local source of GABA. Future studies may reveal the neuronal connectivity of these distinct local CR^+^ interneurons and other roles in the LV-SVZ cell proliferation and neurogenesis processes.

The subep-ChAT^+^ neurons predominantly reside in the ventral domain of the SVZ. I*n vivo* optogenetics stimulation and inhibition of the defined (Cg1-subep-ChAT^+^) circuit is sufficient to modulate SVZ neurogenesis surrounding the subep-ChAT^+^ neurons in the ventral area. While this strongly suggests that subep-ChAT^+^ neurons release ACh locally in the ventral SVZ, further research can be directed to understand their axon terminals in the whole SVZ niche. In consequence, i*n vivo* optogenetics modulation of this circuit is also sufficient to regulate the activity of LV NSCs proliferation and cellular division regionally in the ventral areas of LV-SVZ. Future research may explore the molecular mechanisms of subep-ChAT^+^ neuronal regulation of the LV NSCs activity and cellular divisions in V-SVZ and SVZ, respectively.

Taken together, our results uncover a new neural circuit that allows direct cortical regulation of the LV-SVZ neurogenesis. In humans, there is evidence for active neurogenesis at the wall of the lateral ventricles that generate migratory neuroblasts for up to two years after birth ^34^, but the circuit mechanism is unknown. The analogous process in rodents may shed light on how the neurogenesis during postnatal brain development is influenced by synaptic inputs and neural activity. At present, how cortical inputs to the SVZ are related to environmental factors and behavior remains unclear, but our results promise to bridge neural circuit activity with neurogenesis and open a new avenue of research into cortical activity dependent SVZ neurogenesis in postnatal and adult animals.

## Methods

### Animals

All experiments were approved by the Institutional Animal Care and Use Committee at Duke University. Mice were group housed on a standard 12 h light/dark cycle (lights on at 7 a.m.) with a controlled average ambient temperature of 21 °C and 45% humidity. The following mouse lines were purchased from JAX: *C57BL/6J* (000664); *VGlut1-Cre* (023527); *ChAT-eGFP* (007902); ChR2*(RCL-hChR2(H134R)/tdT)-D (Ai27)* (012567); *ChAT-Cre* (006410); *Cr-Cre* (010774); *VGat-Cre* (016962); *Cb-Cre* (028532); *SST-Cre* (013044); *PV-Cre* (017320); *Ai9(RCL-tdT)* (007905). We generated the *R26R-FLEX-TVA-2A-RabiesG-2A-tdTomato-FLEX* (*R26F-RTT*) mouse line for rabies monosynaptic circuit tracing **(Fig. 2A)**.

### Rabies virus retrograde tracing

An intraventricular approach via a Cre-dependent viral strategy was employed to avoid labeling striatal ChAT^+^ neurons. In the beginning we used a monosynaptic circuit tracing with Rabies strategy^35^ by injecting the first virus into LV of *ChAT-Cre* mice to express TVA, Rabies-G protein, and tdTomato in subep-ChAT^+^ neurons. The infected subep-ChAT^+^ neurons were later targeted with the second EnvA G-deleted rabies virus to enable efficient mono-synaptic tracing. It was extremely difficult to infect the same subep-ChAT^+^ neurons via two separated viral injections. To overcome this obstacle, *R26R-FLEX-TVA-2A-RabiesG-2A-tdTomato-FLEX (R26F-RTT)* mice was successfully generated and validated **(Fig. 2A and Extended Figs. 1, 2 & 3)**. These mice allow for monosynaptic tracing of Cre-targeted neurons via a single EnvA G-deleted Rabies-eGFP (Salk, USA) or EnvA rVSV-eGFP (EnvA/RABVG-eGFP) (Salk, USA) viral injections.

### Stereotaxic injections

Stereotaxic injections were performed as mice were kept deeply anesthetized in a stereotaxic frame (David Kopf Instruments) with isoflurane. For circuit tracing, rabies virus, 300 nL of (EnvA G-deleted Rabies-eGFP or EnvA rVSV-eGFP (Salk, USA)) virus were injected into striatum or lateral ventricle (LV) of P30 *ChAT-Cre; R26F-RTT* mice. Viruses were infused slowly over 10 min into striatum as the following coordinates relative to Bregma (AP: +1, ML ± 2.0, DV: 2.2 from brain surface) or LV (AP: +0.8, ML ± 0.65, DV: 2.1 from brain surface) using a microdriver with a 10 μL Hamilton syringe. For electrophysiological testing of Cg1 inputs to the subep-ChAT^+^ neurons (optogenetic-light stimulation), adeno-associated viral viruses were used for Cre-dependent expression of the excitatory channelrhodopsin. pAAV-CaMKIIa-hChR2(E123A)-mCherry (addgene #35506) (300nl) virus or pAAV-CaMKIIa-hChR2(E123T/T159C)-EYFP (addgene #35509) (300nl) virus were injected into P30 *C57BL/6J* or P30 *ChAT-Cre; R26F-RTT* mice, respectively. Also, pAAV-Ef1a-DIO hChR2(E123T/T159C)-mCherry (addgene #35510) (300nl) virus was injected into P30 *VGlut1-Cre; ChAT-eGFP* and P28 *Cr-Cre; ChAT-eGFP* mice. Viruses were infused slowly over 10 min into the Cg1 using the following coordinates relative to Bregma (AP: +0.8, ML ± 0.3, DV: 0.5 from brain surface). For pAAV retrograde tracing, pAAVrg-hSyn-DIO-mCherry (addgene #50459-AAVrg) and pAAVrg-hSyn-mCherry (addgene # 114472-AAVrg) viruses (300 nl) was injected into the LV P28 *Cr-Cre*, P30 *C57BL/6J* (P30) and P30 *VGlut1-Cre*. Viruses were infused slowly over 10 min into the Cg1 as the following coordinates relative to Bregma (AP: +0.8, ML ± 0.65, DV: 2.1 from brain surface) .For *in vivo* optogenetic testing of Cg1 inputs to the subep-ChAT^+^ neurons, pAAV-Ef1a-DIO hChR2(E123T/T159C)-mCherry (addgene #35510) (300nl) and pAAV_hSyn1-SIO-stGtACR2-FusionRed (addgene #105677) (300nl) viruses were injected into P28 *Cr-Cre* mice. Viruses were infused slowly over 10 min into the Cg1 (AP: +0.8, ML ± 0.25, DV: 0.5 from brain surface).

All viruses were infused slowly for over 10 min using a Nanoject (Drummond Scientific) connected to a glass pipette. The injection pipette was left in place for 10 min post-injection before it was retracted.

### SVZ wholemount preparation and whole-cell patch-clamp recording

For electrophysiology experiments both male and female mice (4-to 10-week-old) were used. They were anesthetized with isofluorane, transcardially perfused and then ventricular wall were dissected as SVZ wholemounts in ice-cold NMDG artificial cerebrospinal fluid (ACSF; containing 92 mM NMDG, 2.5 mM KCl, 1.2 mM NaH2PO4, 30 mM NaHCO3, 20 mM HEPES, 2 mM glucose, 5 mM sodium ascorbate, 2 mM thiourea, 3 mM sodium pyruvate, 10 mM MgSO4, 0.5 mM CaCl2), and bubbled with 5% CO2/95% O2. Tissues were then bubbled in same solution at 37 °C for 15 min, transferred to bubbled, modified-HEPES ACSF at 23–25 °C (92 mM NaCl, 2.5 mM KCl, 1.2 mM NaH2PO4, 30 mM NaHCO3, 20 mM HEPES, 2 mM glucose, 5 mM sodium ascorbate, 2 mM thiourea, 3 mM sodium pyruvate, 2 mM MgSO4, 2 mM CaCl2) for at least 45 min before start experimentation. (For NMDG artificial cerebrospinal fluid and modified-HEPES ACSF solutions, pH was adjusted to 7.5 and osmolarity to 290 mOsm). Recordings were performed in submerged chamber, superfused with continuously bubbled ACSF (125 mM NaCl, 2.5 mM KCl, 1.25 mM NaH2PO4, 26 mM NaHCO3, 20 mM glucose, 2 mM CaCl2, 1.3 mM MgCl2) at 2.5–5 ml/min at 23–25 °C.

Patch electrodes with a resistance of 3–6 MΩ were pulled from borosilicate glass capillaries using a horizontal puller (P-97, Sutter-instruments). All subep-ChAT^+^ neurons used in the electrophysiology recordings were identified by their distinctive morphology from striatal cholinergic neurons, location (within 15-20 µM from LV surface), their non-spontaneous firing activity, and their depolarized resting membrane potentials (∼-45 mV). For measuring EPSCs when holding at – 60 mV, the following internal solution was used: 130 mM potassium gluconate, 2 mM NaCl, 4 mM MgCl2, 20 mM HEPES, 4 mM Na2ATP, 0.4 mM NaGTP and 0.5 mM EGTA, pH adjusted to 7.2 with KOH. A 40 µM picrotoxin ((124-87-8), Millipore Sigma) was added in the bath solution in voltage-clamp mode. For measuring inward IPSCs when holding at – 60 mV, we used internal solution containing K-Cl (high Chloride) KCl (135 Mm), HEPES (10 mM), Na2ATP (2mM), NaGTP (0.2 mM), MgCl2 (2 mM), and EGTA (0.1 mM), pH adjusted to 7.2 with KOH. To block glutamatergic transmission, 50 µM DL-AP5 (NMDA antagonist) ((3693/50), R&D Systems) and 50 µM CNQX (AMPA antagonist) ((0190), Tocris) were added in the bath solution in voltage-clamp mode. To block action potentials in the patched cell, the sodium channel blocker QX-314 (10 mM, 552233, Millipore Sigma) was also added. Signals were amplified with Multiclamp 700B (filtered at 10 kHz), digitized with Digidata 1440A (20 kHz), and recorded using pClamp 10 software (Axon). Light-activation of channelrhodopsin was achieved using a 473-nm laser (X-Cite exacte) through a 40x objective (Nikon). Action potentials, evoked EPSCs and evoked IPSCs were analyzed using AxoGrapth (AxoGrapth). One or two subep-ChAT^+^ neurons were recorded per wholemount SVZ. Up to three neurons from each animal were recorded. Measurements were taken as an average of at least five responses to obtain a data point.

For focal (targeted) photostimulation of local CR^+^-IPSCs from subep-ChAT^+^ neurons, we used Polygon 400 Digital Micromirror Device (Mightex; multiwavelength pattered illuminator) to control the temporal dynamics (the size, shape, intensity, and position) of light inputs. The illumination consisted of a 100 ms light pulse (470-nm) at 50% intensity in the selected area (50 µm). The pulse was triggered using a TTL pulse from the Digidata to synchronize the stimulation with electrophysiology. Photostimulation was performed for one fluorescent subep-CR^+^ neuron presents in the field of view resulted in postsynaptic effects in the recorded neuron.

### Immunofluorescence staining and imaging

Preparation of brain tissue for IHC staining was described previously ^25^. We used primary antibodies to GFP (#GFP-1020, 1:400, AVES lab), Choline Acetyltransferase Antibody (#AB144P, 1:250, Millipore sigma), tdTomato [16D7] (#EST203, 1:200, Kerafast), RFP (#600-401-379, 1:250, Rockland), Calretinin (#ab702, 1:200, Abcam), calretinin (#6B3, 1:250, Swant), Calretinin (#MCA-3G9, 1:250, EnCor Biotechnology), RFP (#ab62341, 1:200, Abcam), Doublecortin (#AB2253, 1:250, Millipore), Doublecortin (#4604S, 1:200, Cell Signaling Technology), Doublecortin (#MA5-17066, 1:200, ThermoFisher Scientific), EGFR (phospho Y1068) (#ab5644, 1:250, Abcam), EGFR (phospho Y1068) [Y38] (#ab32430, 1:250, Abcam), GFAP (#GFAP, 1:500, Aves Labs). In brief, for circuit tracing and *in vivo* optogenetic experiments, after experiments mice were deeply anesthetized with isoflurane, perfused transcardially with phosphate buffered saline (PBS), followed by 4% PFA in PBS. The perfused brains were removed and postfixed overnight at 4°C in 4% PFA. The fixed brains were either cut into 50 μm coronal sections by Precisionary Instruments VF-500-0Z vibrating microtome or the SVZ wholemounts were dissected. The coronal slices or SVZ wholemounts were incubated in a blocking solution containing 5% donkey serum and TBST for 100 min at room temperature. The sections were then incubated at room temperature overnight in PBS containing 1% donkey serum and antibodies. They were then washed with PBS and incubated with secondary antibodies, Alexa-594 (1:1000, LifeTech) or Alexa-488 (1:1000, LifeTech) or Alexa-647 (1:1000, LifeTech) for 2 h at room temperature, before washing with PBS. The stained sections were counterstained with a 4′,6-diamidino-2-phenylindole solution (DAPI; (D9542) Sigma-Aldrich). After washing four times with TBST, the sections were coverslipped with Fluoromount (Sigma) aqueous mounting medium. Images were taken using Leica SP8 upright confocal microscope (Zeiss) with 10 ×, 20 × and 40 × objectives under the control of Zen software (Zeiss). All antibodies used were validated as in previous publications ^25,36^ or by publications available on vendor websites specific to each antibody.

### *In vivo* optogenetic stimulation and inhibition

Cannula targeting the Cg1 region was implanted (AP: +0.9, ML ± 0.25, DV: 0.4 from brain surface) after injecting pAAV-Ef1a-DIO hChR2(E123T/T159C)-mCherry and pAAV_hSyn1-SIO-stGtACR2-FusionRed viruses as described in **Fig. 6A, 7A & 8A**, using implantable mono fiber-optic fiber (200 μm, 0.22 NA, Doric). Protruding ferrule end of cannula was then connected via fiber cord and a rotary coupling joint (Doric) was used to permit free movement. Three to four weeks after viral infection, light-stimulation was delivered by TTL control (Master 8, AMPI) of a 473-nm laser (IkeCool). For *In vivo* optogenetic stimulation, the ipsilateral Cg1 region was stimulated for 10 ms pulses (7-9 mW laser power) at 10 Hz, lasting 10 s, given once every 1 min. For *In vivo* optogenetic inhibition, the Cg1 regions in ipsilateral and contralateral sides were continuously stimulated using 3 mW laser power only to study and avoid overheating in the local cortical areas.

For ChAT, DCX, EGFR (phospho Y1068) [Y38], GFAP, CR and RFP analyses, 50-μm brain coronal sections were cut and collected by Precisionary Instruments VF-500-0Z vibrating microtome or SVZ wholemounts were dissected. The sections or areas surrounding the activated/inhibited subep-ChAT^+^ neurons were selected for analyses.

### *in vivo* EdU labeling

The 5-ethynyl-2’-deoxyuridine (EdU) staining was conducted using EdU Cell Proliferation Kit for Imaging (EdU *in vivo* Kits) (baseclick GmbH, Germany) according to the manufacturer’s protocol. EdU was prepared at 50mg/20mL in sterile PBS and used for pulse labeling of adult mice by performing an intraperitoneal (IP) injection of 500 ul dissolved EdU (50mg/kg). The mice were harvested as described in **Fig. 7 and 8**. The intended coronal slices or SVZ wholemounts for EdU labeling were first stained for other antibodies, and then counterstained with DAPI as described in the immunofluorescence staining section. Subsequently, the sections were washed three times with 3% BSA in PBS. Then, incubated for 30 minutes in a reaction cocktail containing Deionized water, Reaction buffer, Catalyst solution, Dye Azide and Buffer additive while protected from light. After the reaction cocktail was removed, sections were washed three times with 3% BSA in PBS. They were then mounted in vectashield mounting media (vector laboratories Inc, Burlingame, CA) and imaged by using Leica SP8 upright confocal microscope (Zeiss) as above. All steps were carried out at room temperature.

### Quantification and Statistical Analysis

All data are expressed as mean ± SEM and all statistical analyses were performed using a GraphPad Prism (version 8) program. Paired and unpaired t-tests were used for analysis of *in vivo* optogenetics stimulation and inhibition studies. p < 0.05 was considered statistically significant.

## Acknowledgements

We thank Scott Soderling and Shawn Je for helpful comments on the manuscript, Transgenic and Knockout Mouse Facility at Duke University for assistance with generation of R26R-FLEX-TVA-2A-RabiesG-2A-tdTomato-FLEX mice, E. Adlaf, J. Erb, B. Asrican, D. Fromme, P. Paez-Gonzalez, K. Abdi, G. Neves, S. Ramamoorthy, J. David and K. Woldemichael for project assistance. This work was supported by NIH grant R01MH105416.

## Contributions

M.M.N., C.T.K. and H.H.Y. conceived the project and participated in research design. M.M.N. performed all experiments and analyzed data. R.R.K. helped with preparation of SVZ wholemounts for electrophysiology experiments. M.M.N. and H.H.Y wrote the paper.

