## Supplementary figures for "Frontal cortical regulation of neurogenesis and cellular proliferation in the ventral subventricular zone"

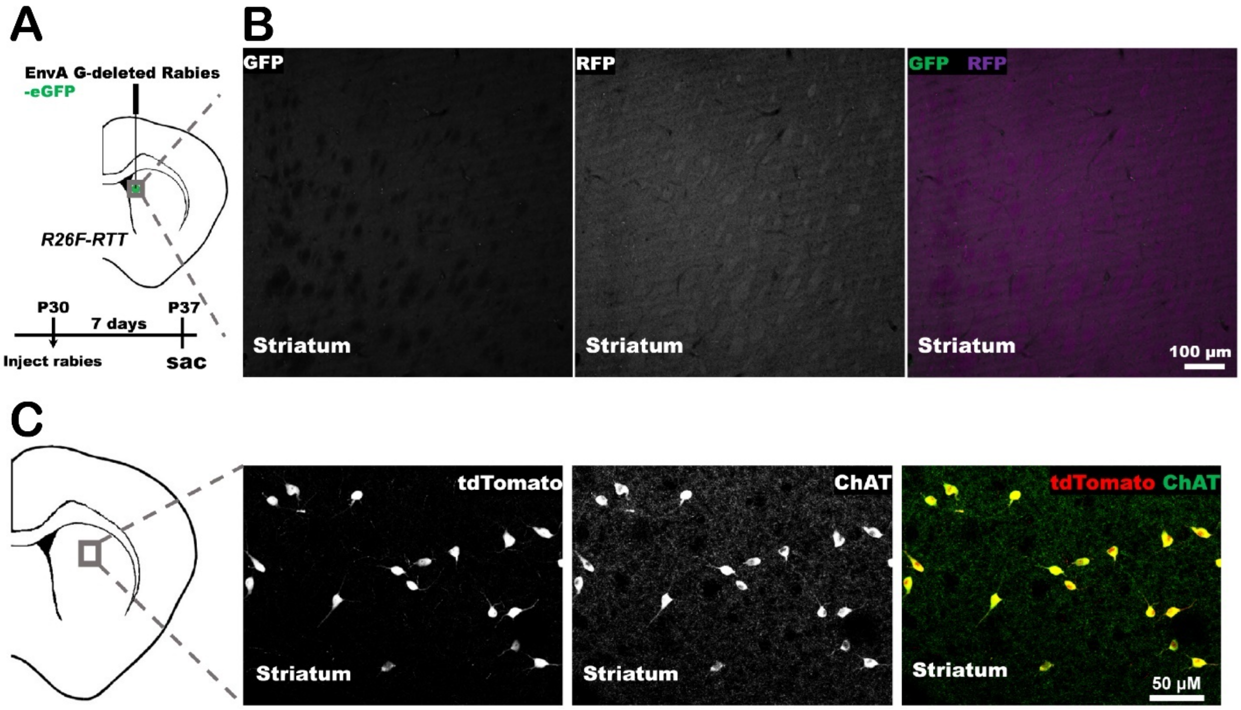


**Extended Figure 1 Validation of *R26F-RTT* mice.**

(A) Schematic representation of EnvA G-deleted Rabies-eGFP virus injection into striatum of P30 *R26F-RTT* mice. Mice were sacrificed 7 days post-injection.

(B) Immunofluorescence staining for GFP (green) and RFP (purple) in striatum of the ipsilateral (injected) side of *R26F-RTT* mice (7 days post-injection).

(C) Immunofluorescence staining for tdTomato (red) and ChAT (green) in striatum region of P30 *Chat-Cre; R26F-RTT* mice.

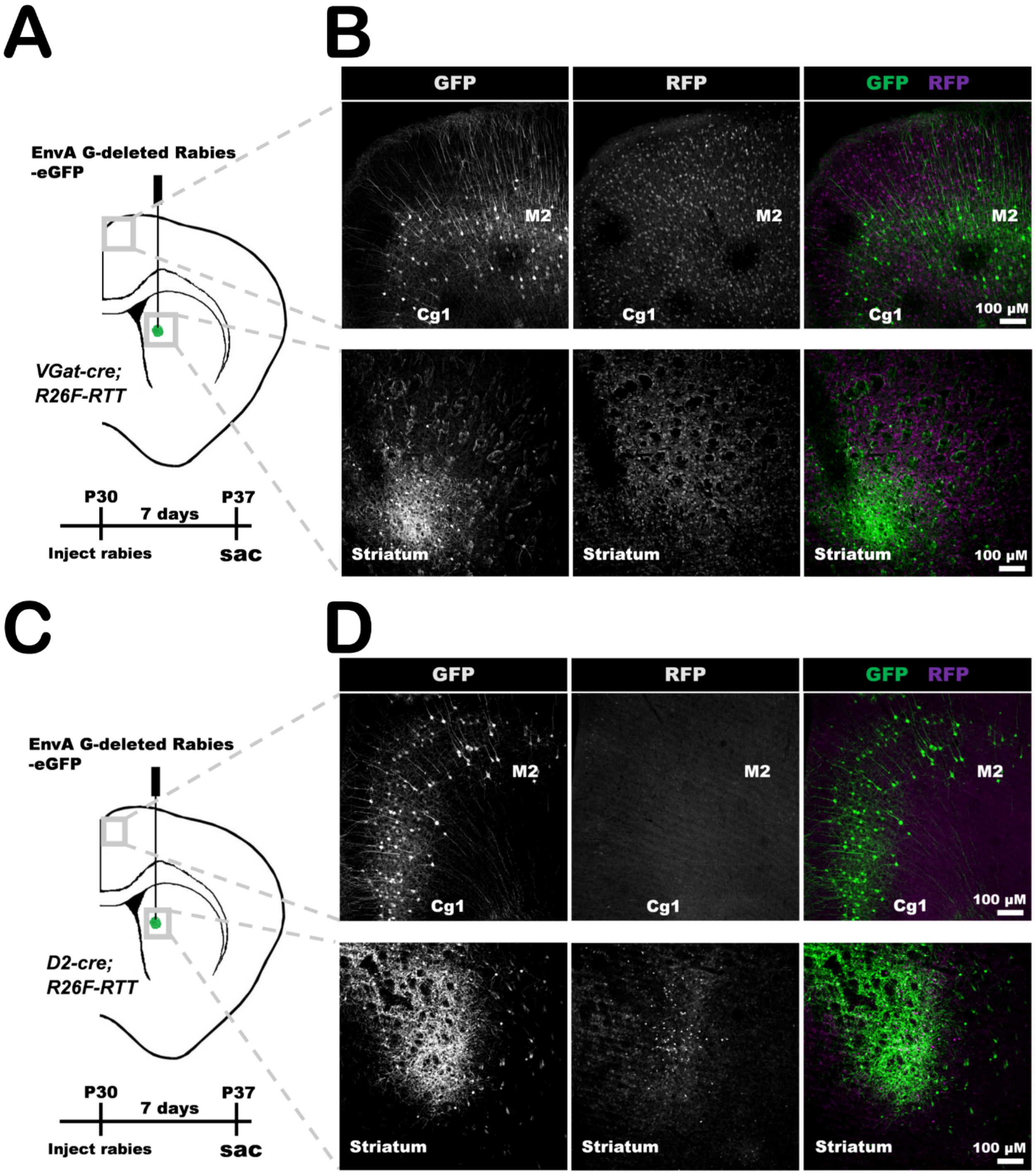


**Extended Figure 2 Rabies injection into the striatum of *VGat-Cre; R26F-RTT* and *D2-Cre; R26F-RTT* mice.**

(A) Schematic representation of EnvA G-deleted Rabies-eGFP virus injection into striatum of P30 *VGat-Cre; R26F-RTT* mice. Mice were sacrificed 7 days post-injection.

(B) Immunofluorescence staining for GFP (green) and RFP (purple) in Cg1 and M2 regions (presynaptic input; upper images), and striatum (injection site; lower images) of *VGat-Cre; R26F-RTT* mice.

(C) Schematic representation of EnvA G-deleted Rabies-eGFP virus injection into striatum of P30 *D2-Cre; R26F-RTT* mice. Mice were sacrificed 7 days post-injection.

(D) Immunofluorescence staining for GFP (green) and RFP (purple) in Cg1 and M2 regions (presynaptic input; upper images), and striatum (injection site; lower images) of *D2-Cre; R26F-RTT* mice.

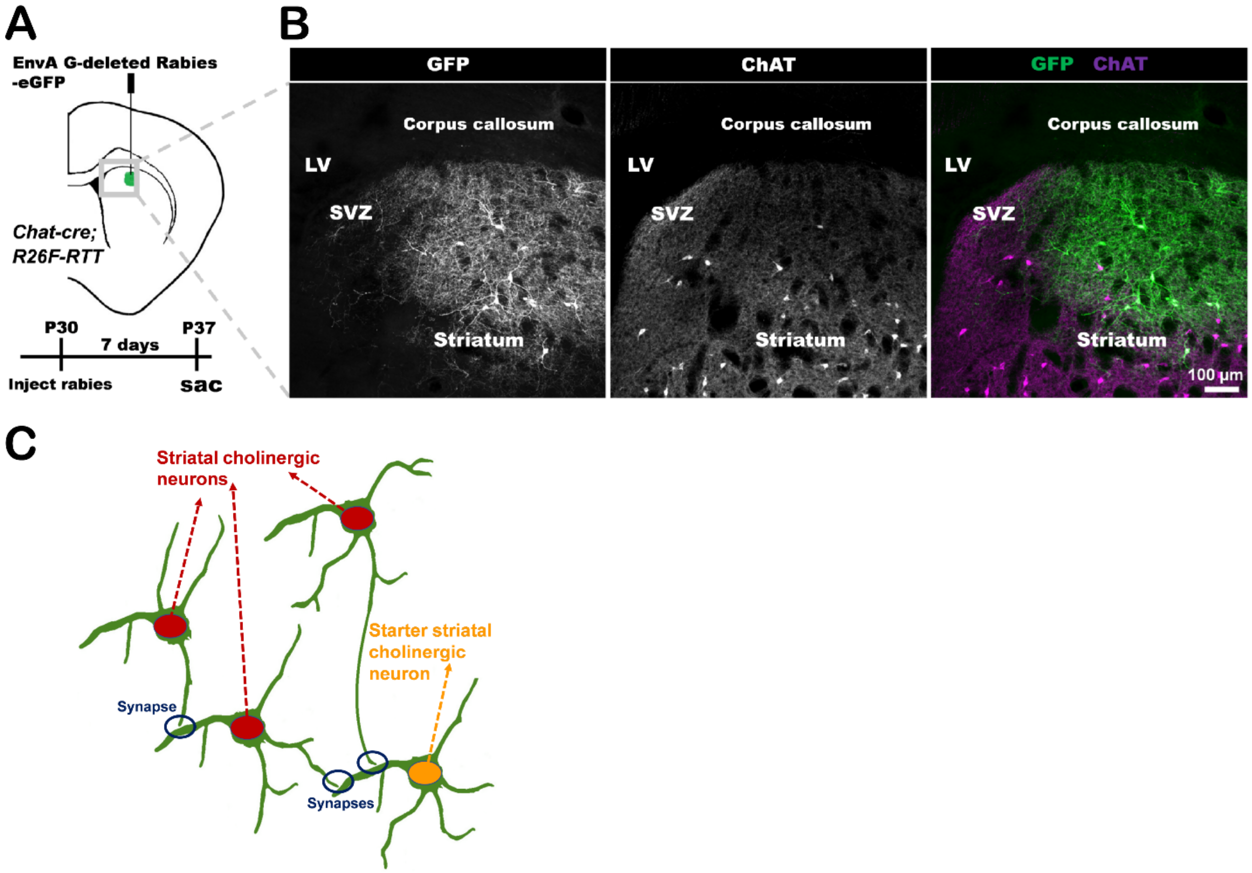


**Extended Figure 3 Rabies injection into the striatum of *chat-Cre; R26F-RTT* mice.**

(A) Schematic of EnvA G-deleted Rabies-eGFP virus injection into striatum of P30 *chat-cre; R26F-RTT* mice. Mice were sacrificed 7 days post-injection.

(B) Immunofluorescence staining for GFP (green) and ChAT (purple) in striatum and SVZ of *chat-cre; R26F-RTT* mice.

(C) Schematic representation of monosynaptic intra cholinergic connections of striatal cholinergic neurons.

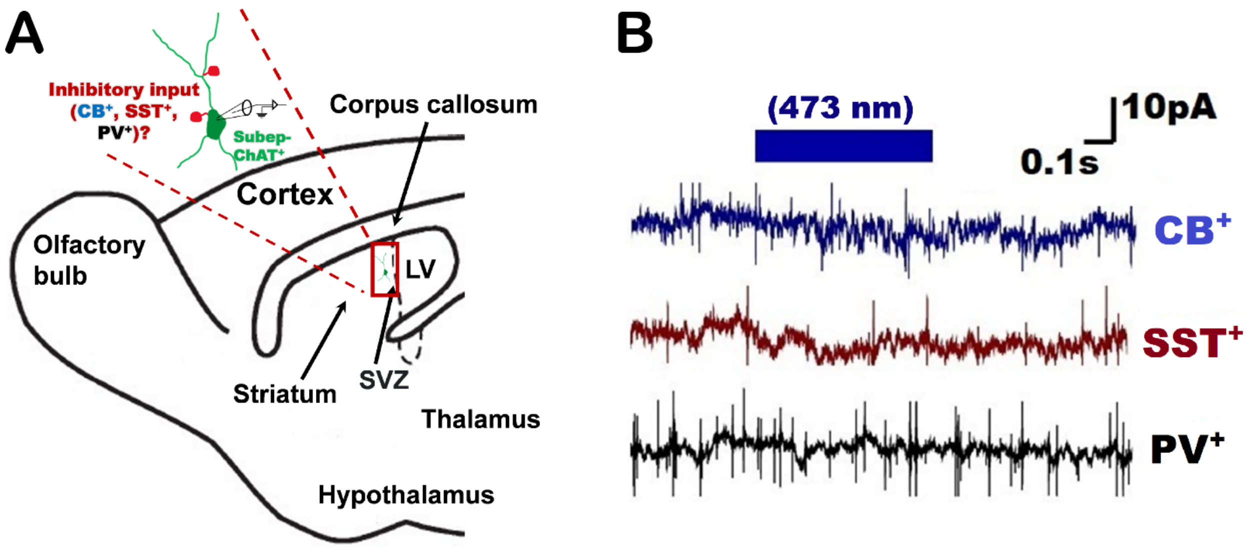


**Extended Figure 4 Other presynaptic inhibitory inputs to the subep-ChAT$^{+}$ neurons.**

(A) Graphical representation for the electrophysiological recording of presynaptic inhibitory inputs into subep-ChAT$^{+}$ neuron of P30-50 *Cb-Cre; ChAT-eGFP; Ai27,* P30-50 *SST-Cre; ChAT-eGFP; Ai27* and P30-50 *PV-Cre; ChAT-eGFP; Ai27* mice.

(B) Representative traces of electrophysiological recordings of IPSCs were obtained in whole-cell voltage-clamp recordings from subep-ChAT$^{+}$ neurons of mice in panel (A) upon 473nm light stimulation for 500ms.

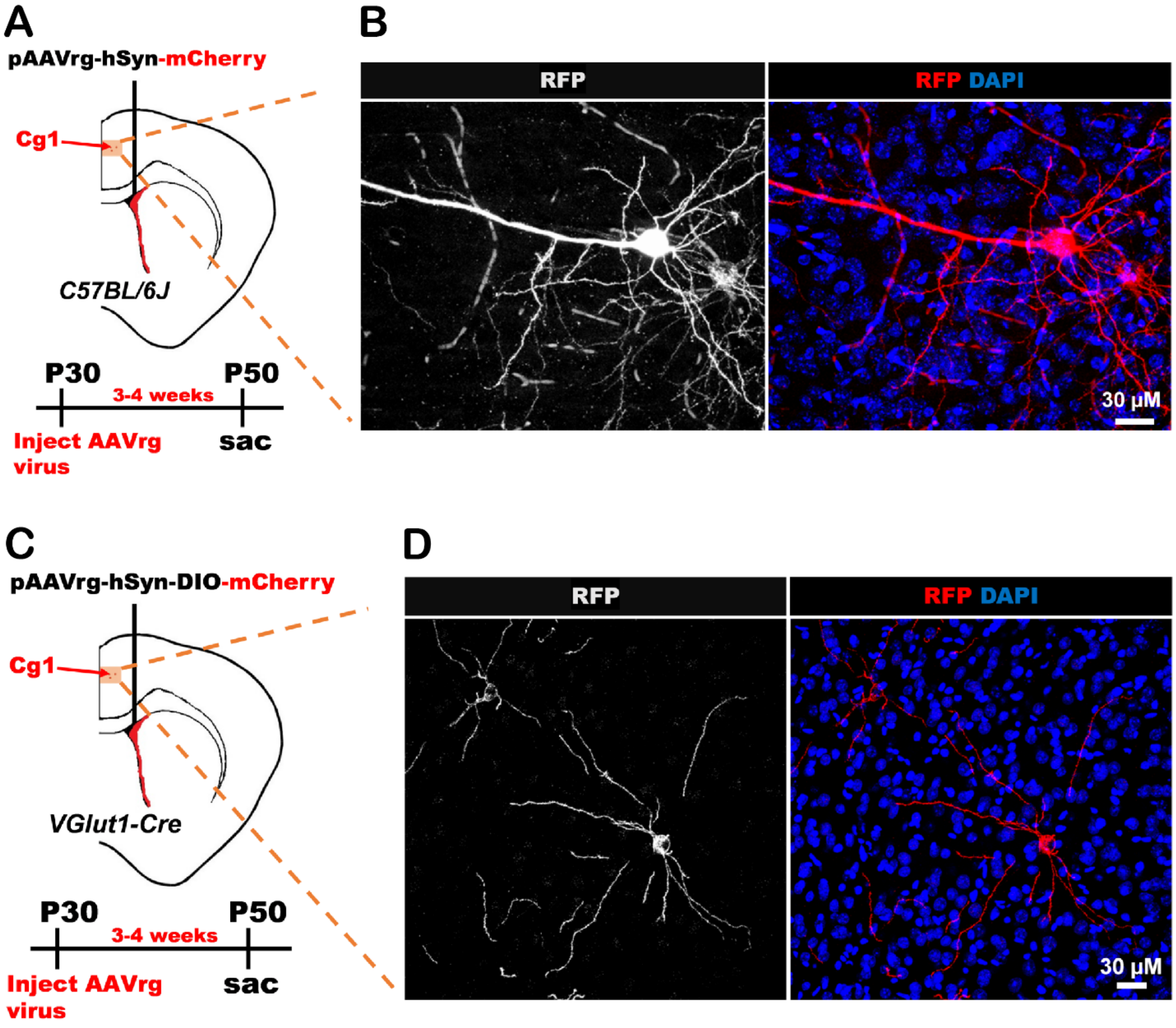


**Extended Figure 5 Tracing neural terminals in the SVZ using pAAV retrograde (pAAVrg) viruses.**

(A) Experimental representation of pAAVrg-hSyn-mCherry virus injection into LV of P30 *C57BL/6J* mice. Mice were sacrificed 3-4 weeks post-injection.

(B) An example of labeled neurons and their projections in ipsilateral Cg1 region. Immunofluorescence staining for RFP (red) in Cg1 region of *C57BL/6J* mice.

(C) Experimental representation of pAAVrg-hSyn-DIO-mCherry virus injection into LV of P30 *VGlut1-Cre* mice. Mice were sacrificed 3-4 weeks post-injection.

(D) An example of labeled neurons and their projections in ipsilateral Cg1 region. Immunofluorescence staining for RFP (red) in Cg1 region of *VGlut1-Cre* mice.

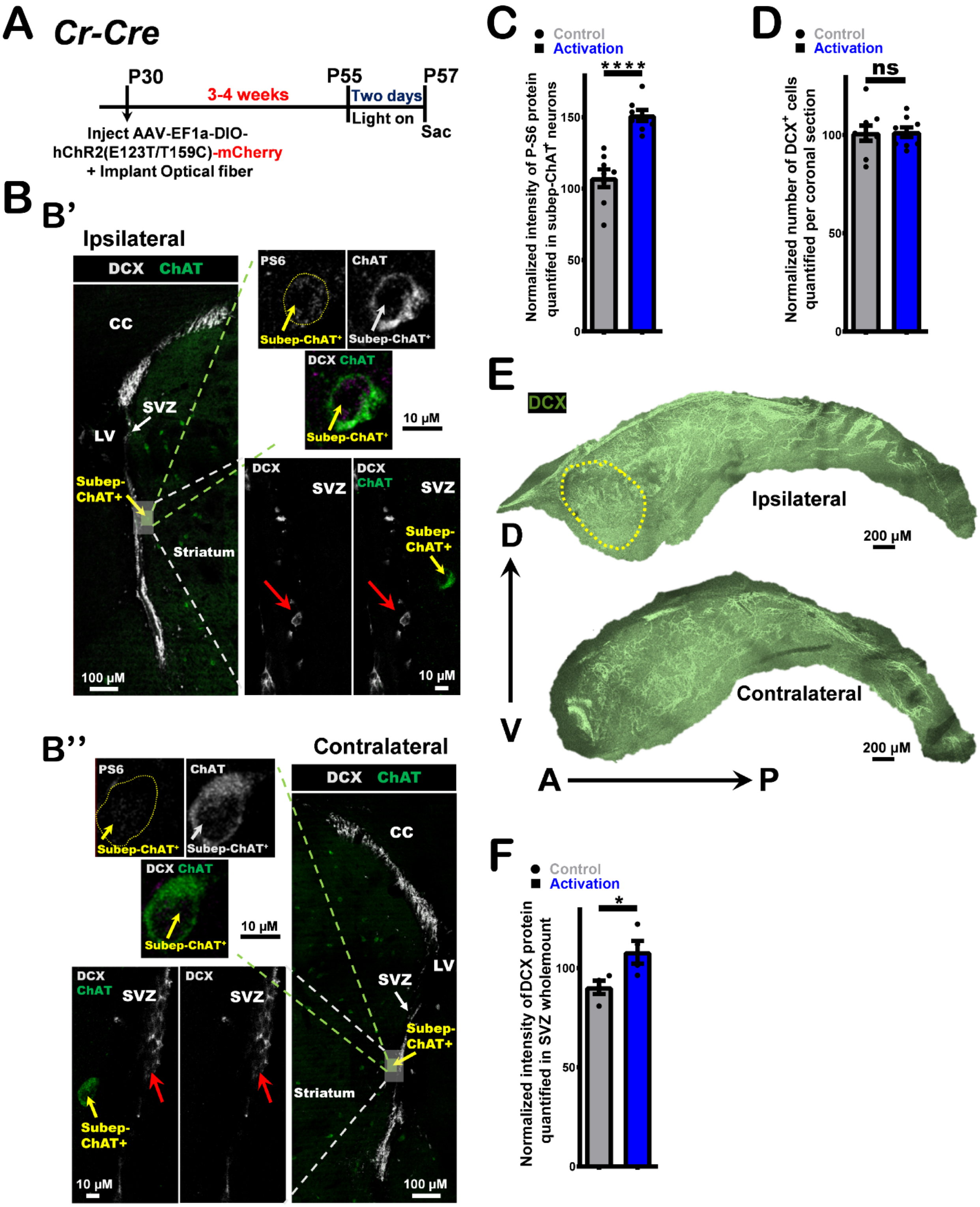


**Extended Figure 6** ***In vivo*** **optogenetic circuit stimulation for two days modulates neurogenesis in the SVZ niche.**

(A) Experimental design of *in vivo* optogenetic stimulation for two days post AAV-EF1a-DIO-hChR2(E123T/T159C)-mCherry virus injection into ipsilateral Cg1 and optical fiber implantation into Cg1 regions of P28 *Cr-Cre* mice

(B) DCX (grey) and ChAT (green) immunofluorescence staining of (B') ipsilateral SVZ wholemount (upper images; activation) vs. (B'') contralateral SVZ wholemount (lower images; control) from stimulated coronal sections of mice in panel A. Identical settings from same brain section were used for imaging. Red arrows represent $DCX^+$ neuroblasts around subep-$ChAT^+$ neurons.

(C) P-S6 intensity analysis of subep-$ChAT^+$ neurons in ipsilateral side (activation) vs. contralateral side (control). P<0.0001, n=8, Paired t-test. Data collected from four stimulated *Cr-Cre* mice. Each dot represents a Subep-$ChAT^+$ neuron.

(D) Analysis of $DCX^+$ neuroblasts in the stimulated coronal sections of ipsilateral side (activation) vs. contralateral side (control). P<0.914 (ns), n=9, Paired t-test. Data collected from four stimulated *Cr-Cre* mice. Each dot represents a total $DCX^+$ cells per coronal section.

(E) DCX (green) immunofluorescence staining of ipsilateral SVZ wholemount (upper images; activation) vs. contralateral SVZ wholemount (lower images; control) from stimulated mice in panel A. Identical settings from same brain wholemounts were used for imaging. Yellow dotted circle represents an area where subep-$ChAT^+$ neurons mainly reside in the ventral region of SVZ.

(F) Analysis of DCX intensity in ipsilateral SVZ wholemount (activation) vs. contralateral SVZ wholemount (control). P = 0.0373, n=4, Paired t-test. Data collected from four stimulated *Cr-Cre* mice. Each dot represents a total DCX protein per SVZ wholemount.

All error bars indicate SEM.

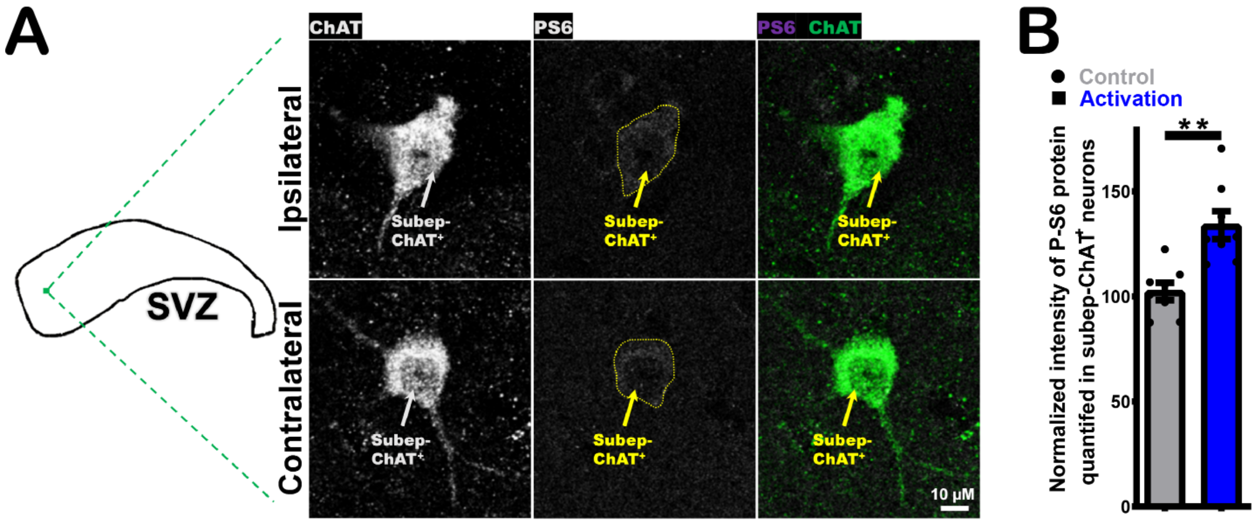


**Extended Figure 7** ***In vivo*** **optogenetic (Cg1-subep-ChAT$^{+}$) circuit stimulation modulates the activity of subep-ChAT$^{+}$ neurons.**

(A) ChAT (green) and P-S6 (purple) immunofluorescence staining of subep-ChAT$^{+}$ neuron in ipsilateral SVZ wholemount (upper images; activation) vs. contralateral SVZ wholemount (lower images; control) from stimulated mice in Fig. 7B. Identical settings from same brain wholemounts were used for imaging.

(B) P-S6 intensity analysis of subep-ChAT$^{+}$ neurons in ipsilateral (activation) vs. contralateral (control) SVZ wholemounts. $P<0.0012$, $n=8$, Unpaired t-test. Data collected from four stimulated *Cr-Cre* mice. Each dot represents a subep-ChAT$^{+}$ neuron.

All error bars indicate SEM.

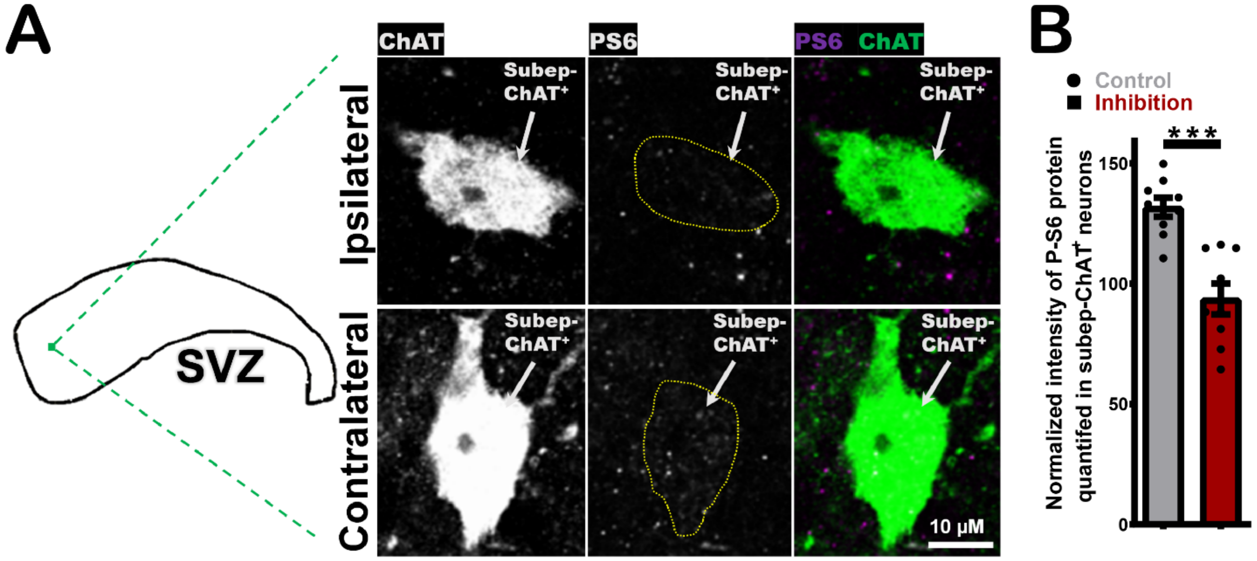


**Extended Figure 8** ***In vivo*** **optogenetic (Cg1-subep-ChAT$^{+}$) circuit inhibition modulates the activity of subep-ChAT$^{+}$ neurons.**

(A) ChAT (green) and P-S6 (purple) immunofluorescence staining of subep-ChAT$^{+}$ neuron in ipsilateral SVZ wholemount (upper images; inhibition) vs. contralateral SVZ wholemount (lower images; control) from stimulated mice in Fig. 8B. Identical settings from same brain wholemounts were used for imaging.

(B) P-S6 intensity analysis of subep-ChAT$^{+}$ neurons in ipsilateral (inhibition) vs. contralateral (control) SVZ wholemounts. P = 0.0001, n=9, Unpaired t-test. Data collected from four stimulated *Cr-Cre* mice. Each dot represents a subep-ChAT$^{+}$ neuron.

All error bars indicate SEM.

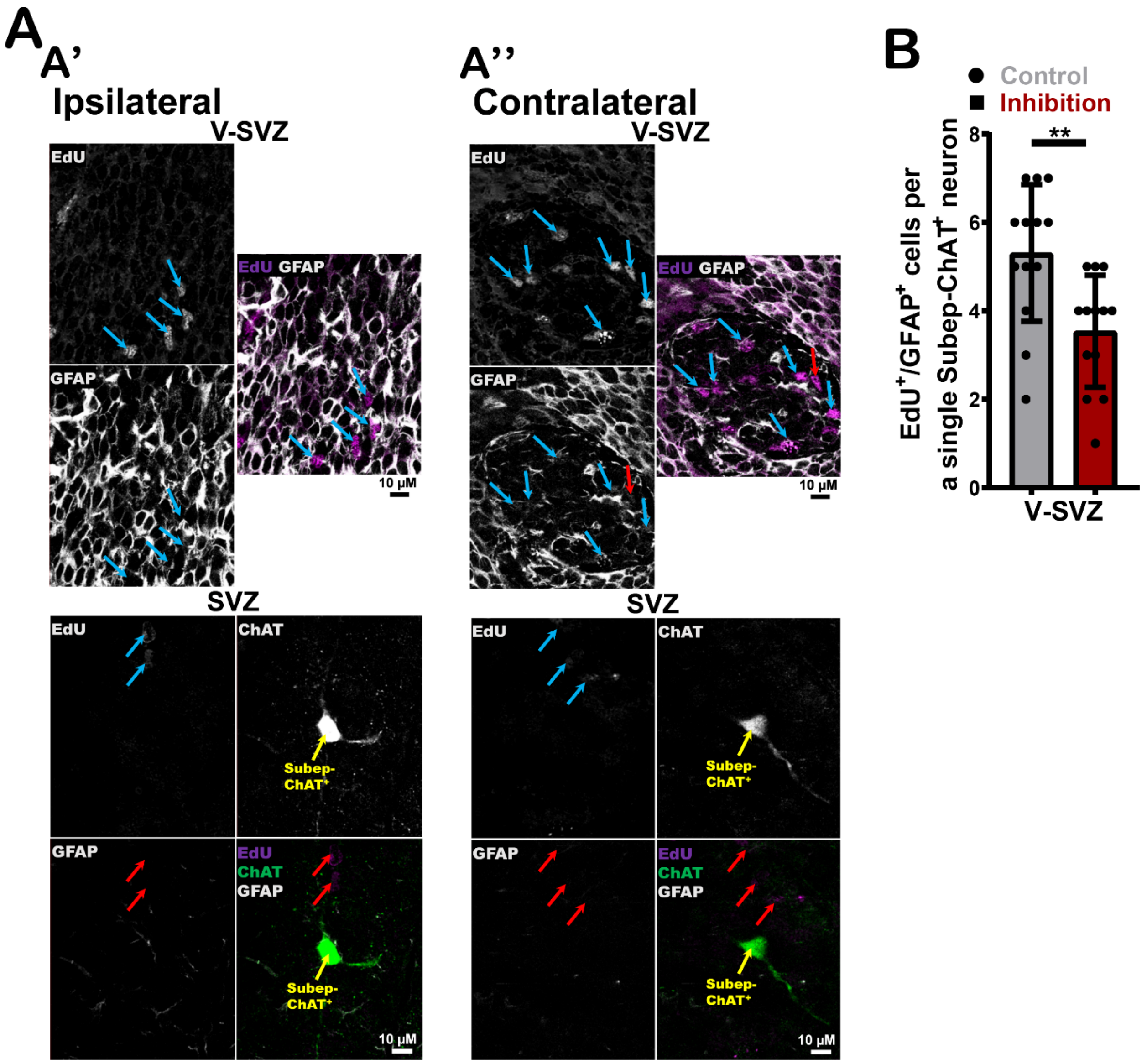


**Extended Figure 9** ***In vivo*** **optogenetic (Cg1-subep-ChAT$^+$) circuit inhibition modulates cellular proliferation in the V-SVZ.**

(A) EdU (purple), ChAT (green) and GFAP (grey) immunofluorescence staining of (A') ipsilateral SVZ wholemount (inhibition) (upper images; V-SVZ and lower images; SVZ) vs. (A'') contralateral SVZ wholemount (control) (upper images; V-SVZ and lower images; SVZ) from stimulated coronal sections of mice in Fig.8B. Identical settings from same brain section were used for imaging. Blue arrows display EdU$^+$/GFAP$^+$ cells surrounding subep-ChAT$^+$ neurons in V-SVZ. Red arrows display EdU$^+$/GFAP$^-$ cells surrounding subep-ChAT$^+$ neurons in V-SVZ and SVZ.

(B) Up: Analysis of EdU$^+$/GFAP$^+$ cells in ipsilateral (inhibition) V-SVZ vs. contralateral (control) V-SVZ of the SVZ wholemounts. P < 0.0048, n=13, Unpaired t-test. Data collected from four stimulated *Cr-Cre* mice. Each dot represents total $EdU^+/GFAP^+$ cells surrounding a subep-$ChAT^+$ neuron.

Down: Analysis of $EdU^+/GFAP^+$ cells in ipsilateral (inhibition) SVZ vs. contralateral (control) SVZ of the SVZ wholemounts. P = 0.0003, n=13, Unpaired t-test. Data collected from four stimulated *Cr-Cre* mice. Each dot represents total $EdU^+/GFAP^+$ cells surrounding a subep-$ChAT^+$ neuron.

All error bars indicate SEM.
